# Early nervous system development in the chaetognath *Spadella cephaloptera* exhibits conserved bilaterian patterning features

**DOI:** 10.64898/2026.03.02.709007

**Authors:** June F. Ordoñez, Alice Frisinghelli, Cristian Camilo Barrera Grijalba, Tim Wollesen

**Author notes:** Corresponding authors Tim Wollesen Email address June F. Ordoñez.

## Abstract

Nervous systems display extensive diversity in structure and organization, yet a broadly conserved set of signaling pathway components and transcription factors is consistently associated with early neurogenesis in many animal lineages. Determining how these conserved markers map onto the spatiotemporal organization of neurogenic territories across phylogenetically informative but underrepresented lineages, particularly within Spiralia, is critical for refining inferences about the evolutionary origins and diversification of nervous systems. Chaetognaths, a spiralian lineage frequently recovered close to Gnathifera, have a compact and centralized nervous system but lack detailed molecular descriptions of early neural development.

Here, we generate an expression-based developmental map of early neurogenesis in the chaetognath *Spadella cephaloptera* by combining nuclear-staining–based anatomical staging with spatiotemporal analyses of conserved developmental genes associated with early neurogenesis and axial patterning from gastrulation through early post-embryonic stages. *Sce-soxB1-like1* and *Sce-neuroD* expressions mark a lateral neuroectodermal territory during gastrulation. Notably, *Sce-neuroD* is activated early in a broad ectodermal domain and is expressed within mitotically active neuroectodermal cells, consistent with early deployment in proliferative neurogenic territories. *Sce-soxB1* and *Sce-soxB2* show delayed and more spatially restricted expression relative to *Sce-soxB1-like1*, suggesting a paralog-specific partitioning of SoxB deployment during chaetognath neurogenesis. *Sce-bmp2/4* and *Sce-chd* exhibit reciprocal dorsoventral expression during gastrulation that coincides with early neurogenic territory formation, before transitioning to more localized expression later in development. *Sce-nk6* and *Sce-hb9* reveal early ventral regionalization of the developing ventral nerve center (VNC), with *Sce-hb9* occupying a subset of a broader *Sce-nk6* domain, in line with conserved ventral subtype-associated regionalization. *Sce-th* (*tyrosine hydroxylase*) is detected in a small bilateral subset of hatchling VNC cells, while *Sce-dbh* (*dopamine beta-hydroxylase*) is first detected only in early juveniles in the anterior VNC and head domains, suggesting stage-dependent and region-specific deployment of catecholamine-pathway components.

Together, these expression-based datasets provide a comparative reference for early neurogenesis in chaetognaths and a framework for assessing conserved and lineage-specific features of early neurogenic patterning across Spiralia.

## Introduction

Nervous systems exhibit remarkable diversity in organization, ranging from basiepithelial nerve nets of cnidarians to the highly centralized brains and nerve cords of many bilaterians (Holland, 2003; Schmidt-Rhaesa, 2007). Despite this architectural divergence, comparative developmental studies consistently identified a shared repertoire of signaling pathways and transcription factors associated with early neural development (Arendt et al., 2016, 2008; De Robertis and Sasai, 1996; Hartenstein and Stollewerk, 2015). In well-studied model systems, signaling pathways (e.g., Bone Morphogenetic Protein (BMP)/Chordin signaling) have been associated with the positioning of neurogenic territories, while transcription factors, most prominently members of the Sox and basic Helix–Loop–Helix (bHLH) families, are widely implicated in neural competence, patterning, and neuronal differentiation (Bertrand et al., 2002; Guth and Wegner, 2008; Kelava et al., 2015; Mizutani and Bier, 2008; Quan and Hassan, 2005; Sprecher and Reichert, 2003; Vervoort and Ledent, 2001). The recurrent involvement of these components across distantly related taxa suggests that early neurogenesis involves deeply conserved molecular signatures (Arendt et al., 2016; Faltine-Gonzalez et al., 2023). However, whether similarities in gene expression correspond to comparable spatial organization of early neurogenic events across lineages remains unclear.

Expanding molecular studies beyond traditional model organisms has begun to refine this picture. Conserved neural-associated transcription factors are widely reported across major bilaterian clades, including acoels, ecdysozoans, deuterostomes, and spiralians (Denes et al., 2007; Janssen et al., 2018; Lowe et al., 2003; Martín-Durán et al., 2018; Martinez et al., 2024). Within Spiralia, studies in annelids, mollusks, and nemerteans demonstrate that many of these genes are expressed during early neurogenesis, yet their timing, spatial relationships, and cellular contexts vary among taxa, and in some cases, even between closely related species (Buresi et al., 2016; Deryckere et al., 2021; Monjo and Romero, 2015a; Simionato et al., 2008; Sur et al., 2020). Such variation underscores that shared molecular markers can be associated with distinct spatiotemporal configurations of neural development. However, available spiralian developmental datasets remain heavily skewed toward lophotrochozoan model taxa, whereas comparable data from Gnathifera, a major spiralian clade sister to Lophotrochozoa, are scarce. This uneven taxonomic sampling constrains assessments of how broadly patterns inferred from lophotrochozoan models generalize across Spiralia and consequently limits broader inferences about conserved features of bilaterian neurogenesis. Moreover, distinguishing deep homology from convergence in nervous system evolution ultimately requires dense comparative sampling, and expression-based maps from understudied spiralian lineages provide an essential baseline (Hejnol and Lowe, 2015).

Chaetognaths (arrow worms) are marine invertebrates frequently recovered as closely affiliated with Gnathifera, based on recent phylogenomic analyses (Laumer et al., 2019; Marlétaz et al., 2019). The adult nervous system has a compact, non-segmented organization, with an anterior cerebral ganglion connected via circumesophageal connectives to an intraepidermal ventral nerve center (VNC) in the trunk (Goto and Yoshida, 1987; Harzsch et al., 2009; Harzsch and Müller, 2007; Rieger et al., 2010). Classical embryological studies described key early features of development, including the ectodermal origin of the VNC (Doncaster, 1902; John, 1933). Subsequent works using immunohistochemical approaches have characterized neuronal subtypes and documented the progressive maturation of neural structures during post-embryonic development (Goto et al., 1992; Perez et al., 2013; Rieger et al., 2011). However, the molecular underpinnings of early neurogenesis in chaetognaths, including the spatial and temporal deployment of genes consistently associated with early nervous system development across diverse lineages, remain largely unexplored, limiting direct comparisons with other spiralians. In this study, we provide an expression-based developmental map of early neurogenesis in the chaetognath *Spadella cephaloptera* by integrating nuclear staining-based anatomical characterization with spatiotemporal analysis of broadly conserved neural-associated genes (including SoxB/bHLH factors such as *soxB1* and *neuroD*, dorsoventral (DV) patterning genes *bmp2/4* and *chordin*, and early subtype-associated markers *nk6* and *hb9*). By tracing gene expression dynamics from gastrulation through early post-embryonic stages, we delineate the spatial and temporal organization of early neurogenic events in this lineage. Together, these data offer an independent comparative reference for evaluating how these markers associated with nervous system assembly are deployed across Spiralia.

## Methods

### Specimen collection and fixation

Field sampling, animal husbandry, developmental staging, fixation and storage procedures described in (Ordoñez and Wollesen, 2025). Briefly, adult *Spadella cephaloptera* were collected from the intertidal zone of Roscoff, France and maintained in aquaria at 14 °C in 35‰ natural seawater at the University of Vienna. Embryos and post-hatching stages (hatchling and early juveniles) were fixed in 4% paraformaldehyde (PFA) prepared in either MOPS buffer or filtered natural seawater, following the buffer composition and fixation times optimized for ISH and HCR in (Ordoñez and Wollesen, 2025). Specimens were washed with PBST (PBS containing 0.1% Tween-20) and stored in methanol at -20 °C until use.

### Candidate gene identification and orthology assessment

Putative homologs of candidate genes were identified from the *S. cephaloptera* draft transcriptome (Wollesen et al., 2023a) through blastx (Altschul et al., 1990) comparisons with sequences from the NCBI non-redundant (nr) protein database. We targeted regulatory genes representing complementary stages of neurogenesis: (i) factors associated with neural competence and early neuroectodermal specification (*soxB1*), (ii) regulators commonly implicated in neuronal differentiation or commitment (*soxB2* and the proneural bHLH *neuroD*), (iii) DV signaling components (*bmp2/4*, *chordin* (*chd*)), and (iv) ventral patterning and motor-program homeobox genes (*nk6*, *hb9*), along with enzymes involved in catecholaminergic neurotransmitter biosynthesis (*dopamine beta-hydroxylase, dbh*; *tyrosine hydroxylase*, *th*).

Protein translations were aligned with MAFFT v7.490 (Katoh, 2005, 2002) implemented in Geneious Prime (v2023.0.1), and trimmed with ClipKit v2.3.3 (Steenwyk et al., 2020). Orthology was assessed with maximum-likelihood (ML) analysis in IQ-TREE v2, employing ModelFinder for site-heterogeneous model selection and 1,000 ultrafast bootstrap and SH-aLRT replicates (Hoang et al., 2018; Kalyaanamoorthy et al., 2017; Minh et al., 2020). Reference bilaterian sequences and the evolutionary model used for each gene are listed in Table S1.

### Whole-mount fluorescent *in situ* hybridization (FISH) and imaging

Gene expression was examined using hybridization chain reactions (HCR-FISH) (Choi et al., 2014, 2018) for all markers except *dbh* and *th*. Experiments followed the protocol of Choi et al. (2018) as modified by Bruce et al. (2021), with additional adjustments from Ordoñez and Wollesen (2025). Specific HCR probe sets were designed using *insitu_probe_generator.py* (https://github.com/rwnull/insitu_probe_generator (Kuehn et al., 2022)) and were synthesized by Integrated DNA Technologies (München, Germany). For *Sce-bmp2/4*, probe concentration was increased threefold because the short transcript length only produced nine pairs of probes.

Due to limited embryo availability, mid-gastrula stage was not assayed for *Sce-soxB1*, *Sce-soxB2*, and selected double-labeling (*Sce-soxB1-lik*e*1*/*neuroD*, *Sce-chd* /*neuroD*, and *Sce-nk6*/*hb9*).

Expression of *dbh* and *th* was detected using traditional chromogenic/fluorescent ISH approach following Ordoñez and Wollesen (2024) and Wollesen et al. (2023b) for riboprobe synthesis, digoxigenin-labeling, and hybridization. These assays were restricted to hatchlings and early juveniles, since the current protocol does not yield reliable signal in embryos (Ordoñez and Wollesen, 2025). All riboprobes, HCR probe sets, and amplifiers are listed in Table S2 and Data Sheet 1.

Images were acquired using Leica TCS SP5 confocal laser scanning microscope (Leica Microsystems, Heidelberg, Germany). All images were processed in Fiji (Schindelin et al., 2012) and Figures were assembled using Inkscape (https://inkscape.org).

## Results

### Orthology assignment of candidate neural regulators

We recovered putative single-copy orthologs of *soxB1*, *soxB2*, *neuroD, bmp2/4*, *chordin*, *nk6*, *hb9*, *th*, and *dbh* from the *Spadella cephaloptera* draft transcriptome. Maximum-likelihood phylogenetic analyses confirmed their identity, with each gene clustering with the corresponding bilaterian orthologs (Additional File 2: Figs. S1–S9).

Aside from *Sce-soxB1*, we identified two highly similar SoxB1-like sequences from the transcriptome, both initially identified as *soxB1* candidates in blastp searches against the nr database. *Sce-soxB1-like1* showed the highest similarity to Sox2-like from *Tubulanus polymorphus* (e-val 2e^-46^; XP_074658127.1), whereas *Sce-soxB1-like2* matched Sox2 from *Trichuris trichiura* (e-val 8e^-44^; CDW55571.1). In the ML phylogeny, the two sequences formed a strongly supported cluster (SH-aLRT = 95.1%; ultrafast bootstrap support = 100%; Fig. S7), suggesting that they represent closely related variants of *soxB1-like* gene in *S. cephaloptera*. Because *soxB1-like1* is the longer and more complete transcript, we used this sequence for HCR probe design to characterize *soxB1-like* expression.

### Anatomical organization of the developing nervous system in embryos of *Spadella cephaloptera*

Classical histological work on *S. cephaloptera* early developmental stages provided the earliest description of its general embryonic territories and cellular organization (John, 1933). Although these observations did not identify neural progenitors or define neurogenic mechanisms (in contemporary terms), they established the basic epithelial architecture in regions that later give rise to neural tissues, such as the brain and VNC. These foundational observations form the basis for our descriptive assessment of early neuroectodermal organization. Here, we complement earlier work by detailing the spatial arrangement and nuclear characteristics (e.g., nuclear shape and DAPI signal intensity) of the presumptive neuroectoderm across the early gastrula, mid-gastrula, late gastrula, and early elongation stages (Fig. 1).

**Figure 1.**
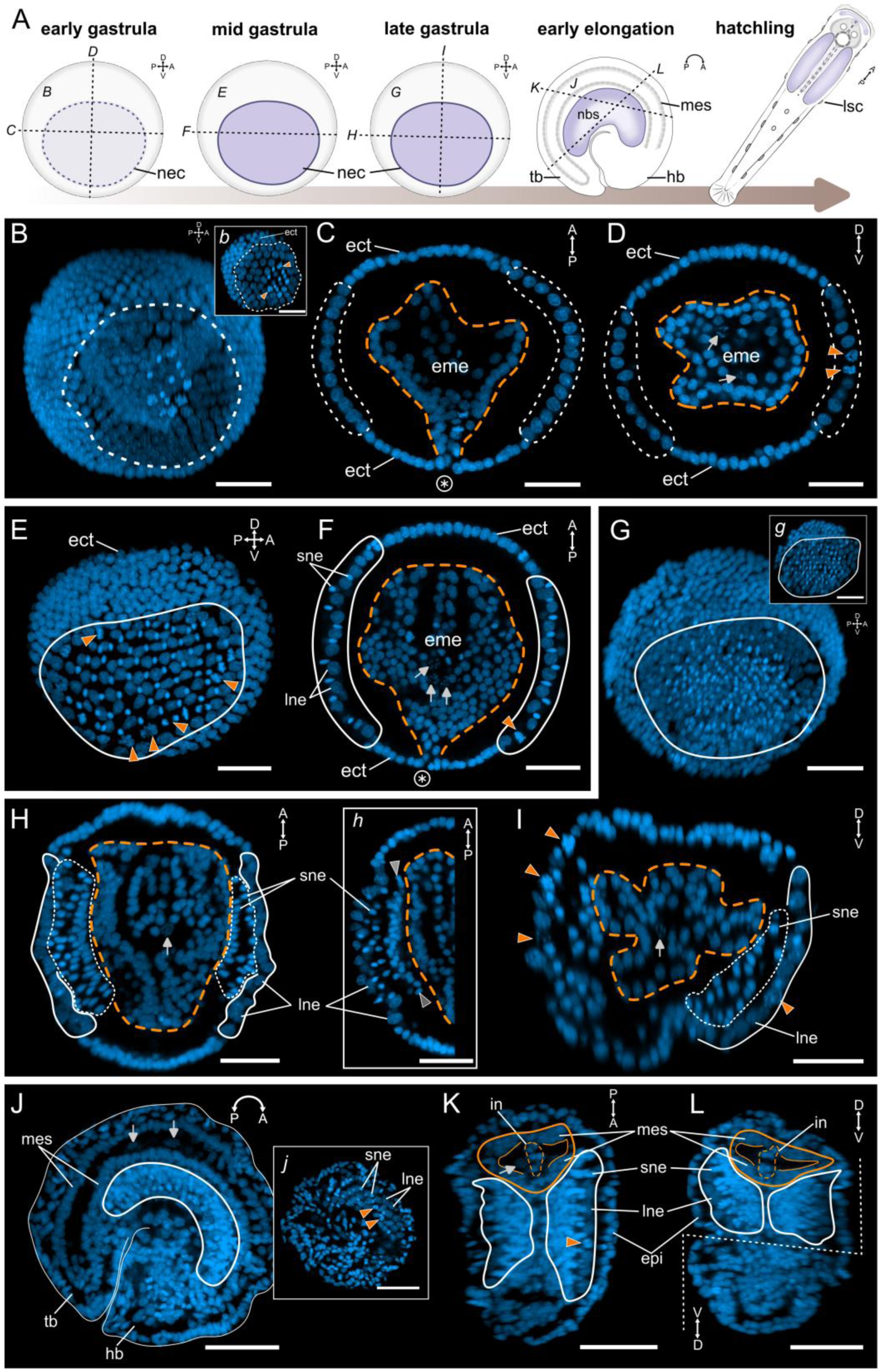
DAPI-based anatomical characterization of ventral nervous system development in *Spadella cephaloptera* embryos. (A) Schematic overview of *S. cephaloptera* development from gastrulation to hatching (early, mid, and late gastrula; early elongation; hatchling); embryos are shown in lateral view and the hatchling in dorsal view. Neurogenic regions, including the neuroectoderm (NEC) and neural cells of the developing ventral nerve cord (VNC), are indicated in purple. At the early gastrula stage, the purple domain corresponds to the presumptive neuroectoderm. (B – D) Early gastrula. (B) Lateral maximum projection. Inset *b* highlights a region of the outer ectoderm showing comparatively large, DAPI-faint nuclei within the presumptive neuroectoderm, contrasted with surrounding non-neural ectodermal nuclei. White dashed outlines demarcate the presumptive neuroectoderm; Orange dashed outlines demarcate the endomesoderm; orange arrowheads indicate morphologically identifiable mitotic figures. (C, D) Dorsal (C) and transverse (D) sections. The encircled asterisk marks the blastoporal region, and gray arrows indicate primordial germ cells. (E, F) Mid gastrula. (E) Lateral maximum projection. (F) Dorsal section. White outlines delineate the NEC. (G – I) Late gastrula. (G) Lateral maximum projection. Inset *g* shows the surface ectoderm highlighting large-nucleus neuroectodermal (LNE) cells and small-nucleus neuroectodermal (SNE) cells within the NEC, as well as surrounding non-neural ectoderm. (H, I) Dorsal (H) and transverse (I) sections showing an outer layer of LNE cells (white outline), an inner layer of SNE cells (dashed white outline), and underlying mesodermal and endodermal tissues (dashed orange outline). Inset *h* highlights the compact inner band of SNE cells at the ectoderm–endomesoderm interface (gray arrowheads). (J – L) Early elongation. (J) Lateral maximum projection. Inset *j* highlights LNE and SNE cells within the nascent ventral nerve cord. (K, L) Dorsal (K) and transverse (L) sections showing the nascent VNC, with the outer LNE layer appear to be positioned subepidermally. Orange outlines demarcate mesodermal cells and the intestinal region (dashed outline). Scale bars: 50 μm. Orientation is indicated in the upper right corner of each panel. Abbreviations: ect, ectoderm; eme, endomesoderm; epi, epidermis; hb, head bud; in, intestine; lne, large-nucleus neuroectodermal cells; mes, mesodermal cells; nbs, neural cells of the developing VNC; nec, neuroectoderm; sne, small-nucleus neuroectodermal cells; tb, tail bud.

During early gastrula, the ectoderm contains a bilateral lateral field of comparatively larger, relatively DAPI-faint nuclei, distinguishable from adjacent epithelia by size (dashed white outlines in Fig. 1A – D). We refer to this field as the presumptive neuroectoderm (NEC), based mainly on its position and its later co-localization with *Sce-neuroD* and *Sce-soxB1-like1* domains described below.

By mid-gastrula, the NEC becomes morphologically recognizable as nuclei within the ventrolateral ectoderm show pronounced size heterogeneity (white outlines in Fig. 1E, F). Cells with large, DAPI-faint nuclei and cells with small, DAPI-bright, and teardrop/ovoid nuclei (visible in transverse and dorsal views) occur in proximity and often alternate along short stretches. For clarity and consistency, we refer to large-nucleus neuroectodermal cells as “LNE cells” and small-nucleus neuroectodermal cells as “SNE cells” throughout the remainder of this manuscript. From mid-gastrula through early elongation, LNE cells within this candidate NEC show frequent mitotic figures (orange arrowheads in Fig. 1), supporting the interpretation that this domain includes a proliferative neuroectodermal population (putative progenitor-like cells).

In the late gastrula, the SNE cells appear to shift basally, populating the ventrolateral margins of the endomesoderm as tightly packed clusters (Fig. 1G – I). Consequently, the ventrolateral ectoderm appears stratified: (i) a surface layer of LNE cells along the outer ectodermal epithelium (white outlines in Fig. 1H, I), (ii) an intermediate subepidermal layer of presumptively delaminating SNE-cells (dashed white outlines in Fig. 1H, I), and (iii) an inner SNE-cell band at the ectoderm–endomesoderm interface (layer between two grey arrowheads in inset *h* of Fig. 1H).

At early elongation, when the embryo already exhibits pronounced C-shaped axial curvature, the inner SNE-cell layer thickens and resolves into bilateral longitudinal bands that converge medioventrally to form the nascent VNC (Fig. 1J – L). LNE cells now appear as a subepithelial layer separated from the epidermis and positioned at the outer margin of the SNE-derived VNC (Fig. 1K, L). In the head bud, John (1933) described a thickened lateral cephalic ectoderm with nuclei positioned nearer the underlying mesodermal masses, interpreted as prospective cerebral ganglion cell precursors. However, in our material, this primordium cannot be delineated with confidence from DAPI morphology alone.

By itself, this nuclear architecture does not define progenitors or precursors, but the staging and terminology (e.g., LNE/SNE cells) described provide the positional framework used in the subsequent sections to map and interpret gene-expression domains.

### Expression of *soxB* paralogs and *neuroD*

#### Sce-soxB1

*Sce-soxB1* expression is not detected in the early gastrula stage (not shown). By the late gastrula stage, *Sce-soxB1* expression appears as bilateral longitudinal strips, predominantly in the inner SNE-cell band (Fig. 2A, B). During early elongation, *Sce-soxB1* is expressed in a subset of cells of the nascent lateral somata clusters (Fig. 2C, D), and later in the presumptive anteroventral cephalic ganglion anlage, from which the vestibular and esophageal ganglia are inferred to arise. In hatchlings, *Sce-soxB1* expression is present in the eyes, a subset of cells surrounding the base of the future mouth, and the lateral somata clusters (Fig. 2E, F; Fig. S9A – G). Weaker *Sce-soxB1* signal is also observed in the cerebral ganglia and medioventral somata clusters (Fig. 2F; Fig. S9D, F).

**Figure 2.**
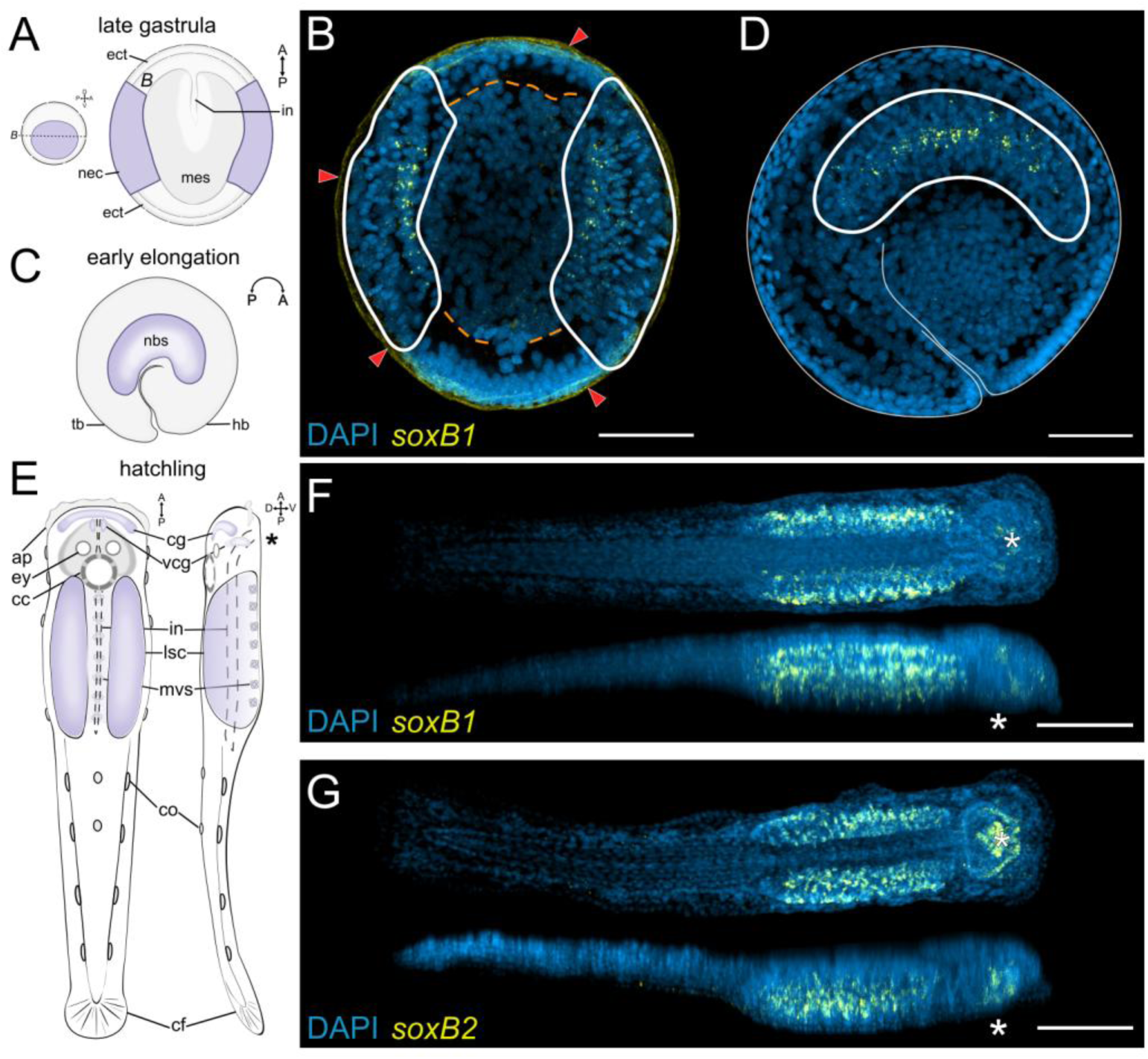
Expression patterns of *Sce-soxB1* and *Sce-soxB2* during embryonic and early post-embryonic development of *Spadella cephaloptera* (A, C, E) Schematic representations of late gastrula (A), early elongation (C), and hatchling (E) stages. The late gastrula is shown in dorsal view, the early elongation stage in lateral view, and the hatchling in dorsal (left) and lateral (right) views. Neurogenic regions, including the NEC and neural cells of the VNC, are indicated in purple. (B, D, F) *Sce-soxB1* expression. (B) Dorsal maximum projection of a late gastrula embryo. White outlines demarcate the NEC, and the dashed orange outline marks the endomesoderm. Red arrowheads indicate non-specific signal from the inner shell layer. (D) Lateral maximum projection of an early elongation embryo in lateral view. (F) Maximum projections of a hatchling, shown in dorsal (top) and lateral (bottom) views. (G) *Sce-soxB2* expression in the hatchling. Maximum projections shown in dorsal (top) and lateral (bottom) views. Scale bars: 50 μm, except (D, E): 100 µm. Orientation is indicated in the schematic representations. The asterisk marks the position of the mouth. Abbreviations: ap, cephalic adhesive papillae; cc, corona cilata; cf, caudal fin; cg, cerebral ganglion; co, ciliary tuft/fence organ; ect, ectoderm; epi, epidermis; ey, eye; hb, head bud; in, intestine; lsc, lateral somata clusters; mes, mesodermal cells; mvs, medioventral somata clusters; nbs, neural cells of the developing VNC; nec, neuroectoderm; tb, tail bud; vcg, anteroventral cephalic ganglion anlage

#### Sce-soxB2

*Sce-soxB2* was not assayed for early and mid gastrula stage individuals and it was not detected in the late gastrula and early elongation stage (not shown). At hatching, *Sce-soxB2* is expressed in the presumptive anteroventral cephalic ganglion anlage, in cells surrounding the future mouth opening, in the presumptive perioral epidermis, and in the lateral somata clusters (Fig. 2G; Fig. S9H – K).

#### Sce-soxB1-like1 *and* Sce-neuroD

Because *Sce-soxB1* is not detectable until late gastrulation and *Sce-soxB2* is detected at hatching, we used *Sce-soxB1-like1* as the primary SoxB marker to map early neurogenic territories and compared it with *Sce-neuroD* using double HCR labeling. In the early gastrula, *Sce-neuroD* is expressed in small bilateral mid-lateral patches within the presumptive neuroectodermal region and *Sce-soxB1-like1* forms a broader lateral domain that overlaps the *Sce-neuroD* center (Fig. 3A – C; Fig. S10D).

**Figure 3.**
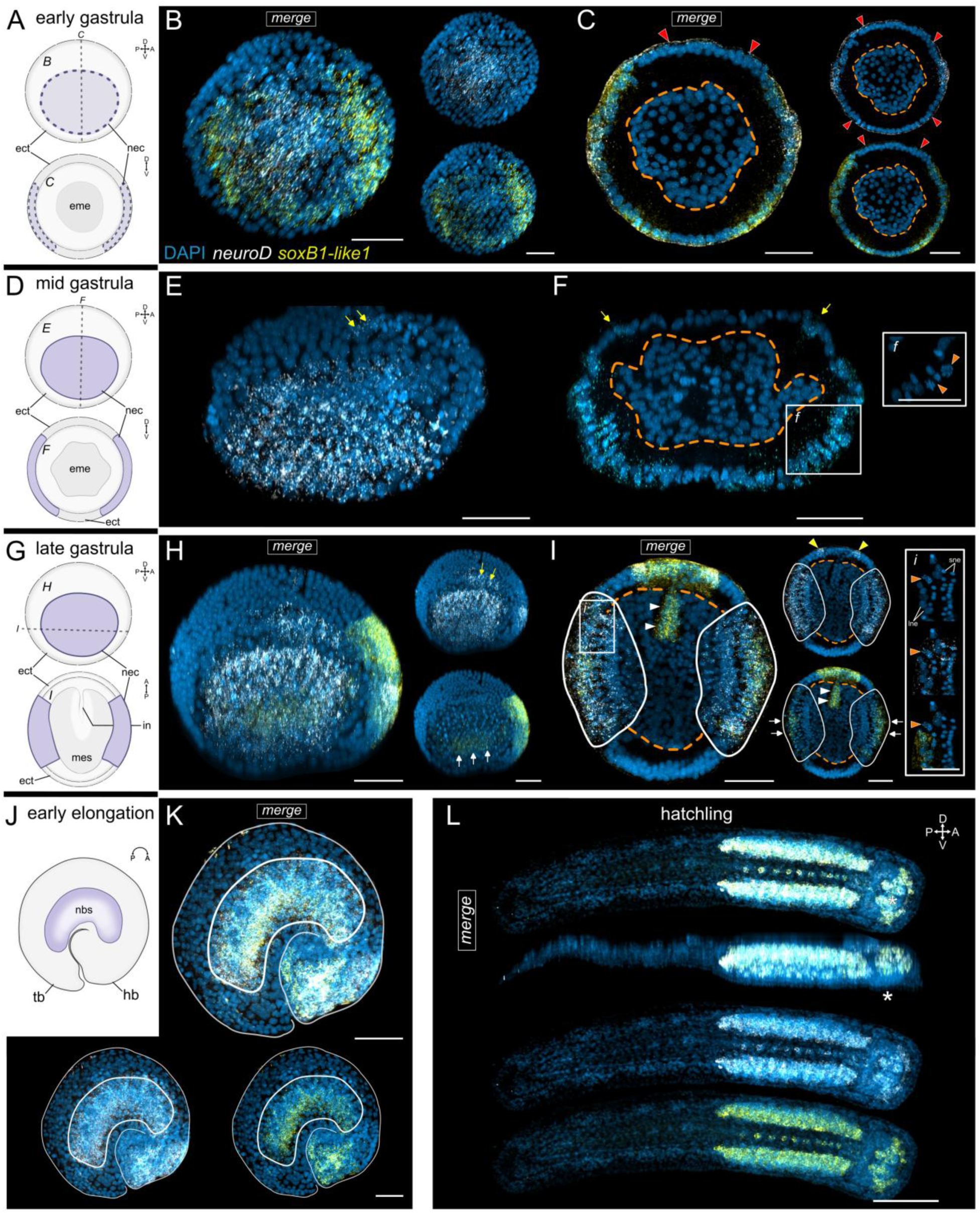
Double-label expression patterns of *Sce-soxB1-like1* and *Sce-neuroD* during embryonic and early post-embryonic development of *Spadella cephaloptera*. Where applicable, fluorescence panels are shown as a merged image (DAPI + probe signals) together with the corresponding single channels (DAPI and each probe). (A, D, G, J) Schematic representations of early gastrula (A), mid gastrula (D), late gastrula (G), and early elongation (J) stages. Early and mid gastrula stages are shown in lateral (top) and transverse (bottom) views; the late gastrula is shown in lateral (top) and dorsal (bottom) views; the early elongation stage is shown in lateral view. Neurogenic regions, including the NEC and neural cells of the nascent VNC, are indicated in purple. (B, C) Early gastrula. (B) Lateral maximum projection. (C) Transverse section. Red arrowheads indicate non-specific signal from the inner shell layer. (E) Mid gastrula. (E) Lateral maximum projection. (F) Transverse section. Yellow arrows indicate expression detected outside the anatomically defined NEC. Inset *f* shows the DAPI staining, with orange arrowheads indicating nuclei exhibiting identifiable mitotic morphology. (H, I) Late gastrula. (H) Lateral maximum projection. Yellow arrows indicate expression detected outside the anatomically defined NEC and white arrows mark the *Sce-soxB1-like1* expression in the ventral NEC. (I) Dorsal section. Yellow arrowheads highlight paired anterior ectodermal *Sce-neuroD* expression domains and white arrowheads mark the anterior endomesoderm domain of *Sce-soxB1-like1*. Inset *i* shows corresponding views of DAPI-only staining (top), *Sce-neuroD* (middle), and *Sce-soxB1-like1* (bottom). Orange arrowheads indicate the same nuclei across all channels, which exhibit identifiable mitotic morphology based on the DAPI signal. (K) Lateral maximum projection of an early elongation stage embryo. (J) Hatchling. Top two rows: merged dorsal and lateral views. Bottom rows: single-channel views. For anatomical context, refer to Fig. 2A’’. The asterisk marks the position of the mouth. Scale bars: 50 μm, except (L): 100 µm. Orientation is indicated in the schematic representations and, where applicable, in the upper right corner of each panel. Abbreviations: ect, ectoderm; eme, endomesoderm; epi, epidermis; hb, head bud; in, intestine; mes, mesodermal cells; nbs, neural cells of the developing VNC; nec, neuroectoderm; tb, tail bud.

At mid gastrula, *Sce-neuroD* is expressed across the morphologically recognizable NEC and a mid-dorsal ectodermal patch that overlies it (yellow arrows in Fig. 3D – F). *Sce-soxB1-like1* was not assayed for this stage.

By late gastrula, *Sce-neuroD* expression remains active across the NEC (Fig. 3G − I) and in the overlying ectodermal patch (yellow arrows in Fig. 3H; Fig. S10E, F). In the mid-anterior ectoderm, *Sce-neuroD* expression also appears as two bilateral patches (yellow arrowheads in Fig. 3I and Fig. S10E). *Sce-soxB1-like1* expression is detected within the NEC but is largely confined to the ventral LNE-cells (white arrows in Fig. 3H, I; Fig. S10F). In the mid-anterior ectoderm, *Sce-soxB1-like1* expression forms a plate-like domain that overlaps with the anterior bilateral *Sce-neuroD* patches (Fig. 3H, I; Fig. S10E). Additionally, a median anterior endomesoderm domain, presumably within endodermal cells, is positive for *Sce-soxB1-like1* (white arrowheads in Fig. 3I; Fig. S10F).

During early elongation, *Sce-neuroD* and *Sce-soxB1-like1* are extensively co-expressed in the nascent lateral somata clusters (Fig. 3J, K; Fig. S10G, H) and in the head bud, including in the presumptive cerebral ganglion precursors (yellow arrowheads in Fig. S10H). The dorsal epidermis of the head bud also shows intense *Sce-neuroD* expression (Fig. 3K; Fig. S10H). In the hatchling, *Sce-neuroD* is expressed in the cerebral ganglion, presumptive anteroventral cephalic ganglion anlage, eyes, corona ciliata, lateral somata clusters, and medioventral somata clusters (Fig. 3L; Fig. S11). *Sce-soxB1-like1* is co-expressed in all these neural domains except the eyes and the corona ciliata (Fig. S11E).

### Expression of *bmp2/4* and *chordin*

In the early gastrula, *Sce-bmp2/4* is expressed in the dorsal ectoderm (Fig. 4A – C; Fig. S12A – C) and in a subset of endomesodermal cells (yellow arrows in Fig. 4C). In contrast, *Sce-chd* expression occupies a broad domain spanning the lateral ectoderm and extending across the ventral region (Fig. 4A – C; Fig. S12A – C). Double HCR of *Sce-chd* and *Sce-neuroD* shows that *Sce-neuroD* expression occurs within the *Sce-chd+* lateral ectoderm, including the presumptive neuroectodermal region (Fig. 4M; Fig. S10B, C).

**Figure 4.**
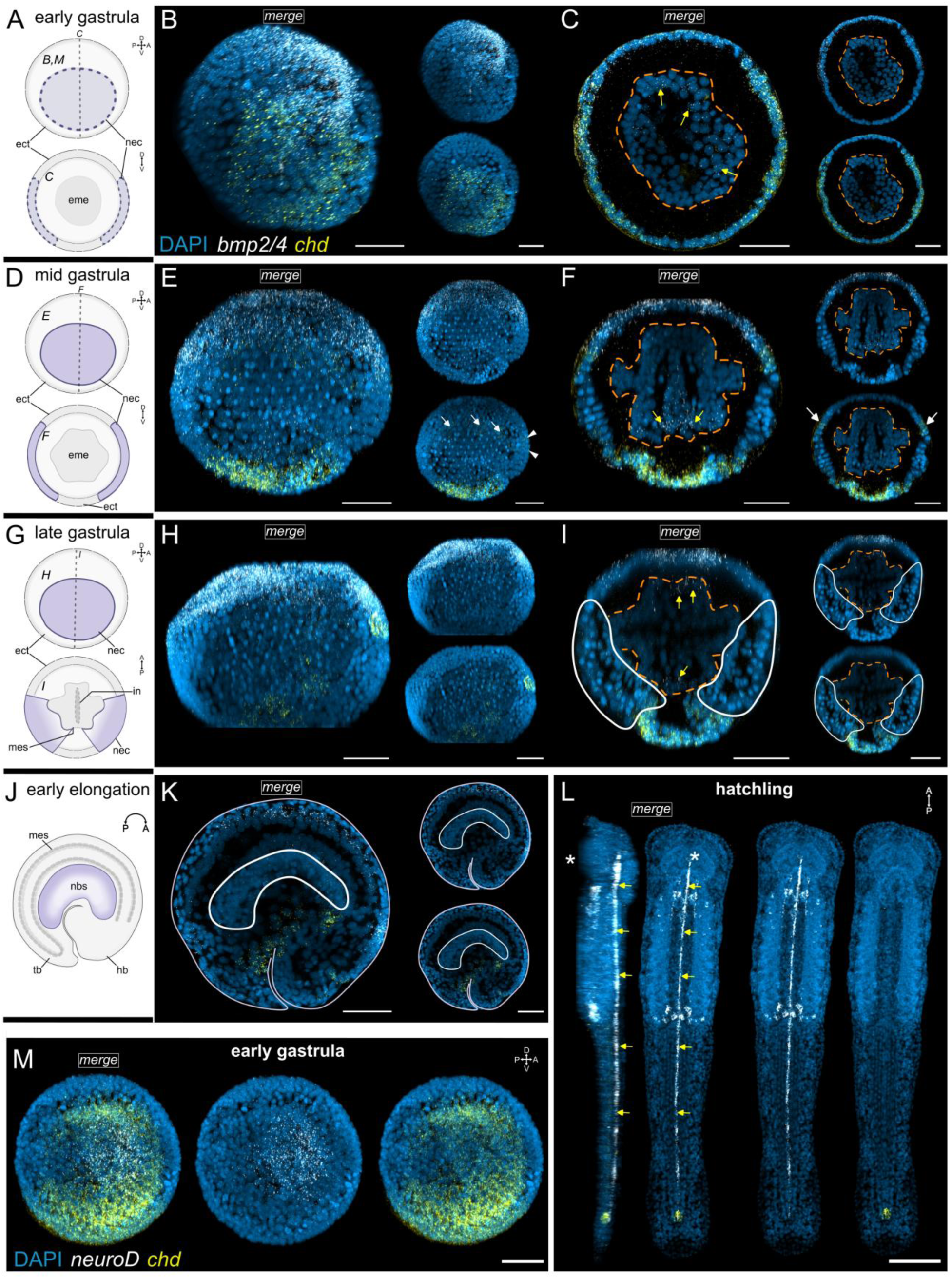
Double-label expression patterns of *Sce-bmp2/4* and *Sce-chd* during embryonic and early post-embryonic development of *Spadella cephaloptera*. Fluorescence panels are shown as a composite overlay (DAPI and two probe signals) together with the corresponding individual probe channels shown with DAPI counterstain. (A, D, G, J) Schematic representations of early gastrula (A), mid gastrula (D), late gastrula (G), and early elongation (J) stages. Gastrula stages are shown in lateral (top) and transverse (bottom) views; the early elongation stage is shown in lateral view. Neurogenic regions, including the NEC and neural cells of the nascent VNC, are indicated in purple. (B, C) Early gastrula. (B) Lateral maximum projection. (C) Transverse section. Yellow arrows (in C, F, and I) indicate *Sce-bmp2/4* expression detected in a subset of endomesodermal cells. Dashed orange outlines (in C, F, and I) demarcate the endomesoderm. (E, F) Mid gastrula. (E) Lateral maximum projection. White arrowheads indicate *Sce-chd* expression in the anterior ectoderm. (F) Transverse section. White arrows indicate *Sce-chd* expression within the dorsal portion of the NEC. (H, I) Late gastrula. (H) Lateral maximum projection. (I) Transverse section. White outlines demarcate the NEC. (K) Lateral section of an early elongation stage embryo. (L) Hatchling. Merged lateral (left) and dorsal (right) views and individual probe channels in dorsal views. Yellow arrows mark the *Sce-bmp2/4* expression in dorsal medial cells. For anatomical context, refer to Fig. 2A’’. The asterisk marks the position of the mouth. (M) Maximum projection of double-label expression of *Sce-neuroD* and *Sce-chd* in an early gastrula embryo shown in lateral view. Scale bars: 50 μm, except (L): 100 µm. Orientation is indicated in the schematic representations and, where applicable, in the upper right corner of each panel. Abbreviations: ect, ectoderm; eme, endomesoderm; epi, epidermis; hb, head bud; in, intestine; mes, mesodermal cells; nbs, neural cells of the developing VNC; nec, neuroectoderm; tb, tail bud.

*Sce-bmp2/4* expression continues to mark the dorsal ectoderm and extends posteriorly towards the blastoporal region in the mid gastrula (Fig. 4D – F; Fig. S12D – G), with persistent expression in some cells of the endomesoderm (yellow arrows in Fig. 4F and Fig. S12G). During this stage, *Sce-chd* expression shows reduced ectodermal signal and is mainly detected in the ventral ectoderm (Fig. 4D – F ; Fig. S12F), with faint expression persisting in the NEC (white arrows in Fig. 4E, F and Fig. S12G), and a low-level signal in the mid-anterior ectoderm (white arrowheads in Fig. 4E and Fig. S12G).

By the late gastrula stage, *Sce-bmp2/4* expression persists in the dorsal ectoderm, extending from the mid-anterior region toward the posterior ectoderm (Fig. 4G – I; Fig. S12H – K). *Sce-bmp2/4* expression also remains detectable in a subset of presumably mesodermal cells (yellow arrows in Fig. 4I). *Sce-chd* expression further weakens in overall intensity and is largely restricted to the ventral ectoderm, with an additional bilateral pair of patches appearing in the mid-anterior ectoderm (Fig. 4G – I; Fig. S12J, K).

In early elongation, *Sce-bmp2/4* is expressed in the dorsal ectoderm of the head bud, in mesodermal cells along the dorsal region of the developing trunk (Fig. 4J, K). *Sce-chd* shows a patchy expression in the ventral and dorsal head bud, the ventral trunk, and tail bud. At hatching, *Sce-bmp2/4* is expressed in the dorsal medial cells forming a continuous longitudinal line that extends from the base of the head to the posterior tail (yellow arrows), in ventral cells of the anterior and posterior boundaries of the trunk, and in a few cells of the lateral somata clusters (Fig. 4L; Fig. S12L – Q). In contrast, *Sce-chd* expression is restricted to a small patch in the posterior tail (Fig. 4L).

### Expression of *hb9* and *nk6*

At the late gastrula stage, *Sce-hb9* is expressed in the presumptive endodermal cells (yellow arrows), whereas *Sce-nk6* expression is broadly detected in the midventral SNE-cell territory (Fig. 5A – C). During early elongation, *Sce-hb9* is expressed in the nascent lateral somata clusters and in the ventral stomodeum (Fig. 5D – F). Expression is also observed in scattered cells of the developing trunk (yellow arrows in Fig. 5E). *Sce-nk6* is similarly expressed in the nascent lateral somata clusters, occupying a more dorsal domain that partially overlaps with *Sce-hb9* expression (Fig. 5E, F). At hatching, *Sce-hb9* is expressed in the presumptive anteroventral cephalic ganglion anlage, perioral epidermis, lateral somata clusters and the gut (Fig. 5G, H; Fig. S13A – D, F – H), while *Sce-nk6* expression is detected in some cells of the head, medioventral somata clusters, the trunk epidermis, and in cells near the trunk–tail boundary (Fig. 5G, H; Fig. S13C – H). Within the lateral somata clusters, *Sce-hb9* domain occupies most of the ventral and mid regions, whereas *Sce-nk6* expression spans a broader domain that includes this *Sce-hb9^-^*dorsal territory (Fig. 5H).

**Figure 5.**
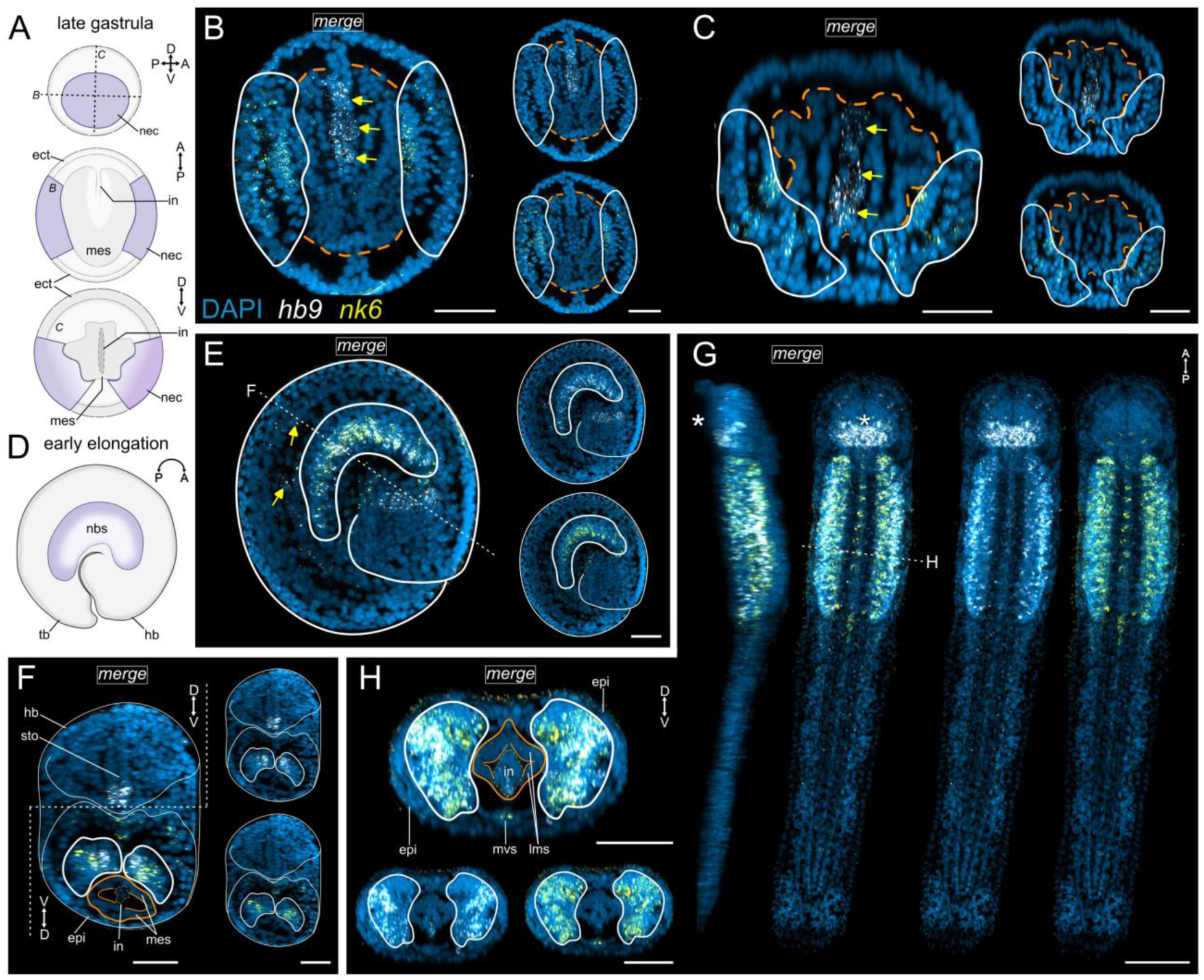
Double-label expression patterns of *Sce-hb9* and *Sce-nk6* during embryonic and early post-embryonic development of *Spadella cephaloptera*. Fluorescence panels are shown as a composite overlay (DAPI and two probe signals) together with the corresponding individual probe channels shown with DAPI counterstain. (A, D) Schematic representations of late gastrula (**A**) and early elongation (**D**) stages. The late gastrula is shown in dorsal (middle) and transverse (bottom) views; the early elongation stage is shown in lateral view. Neurogenic regions, including the NEC and neural cells of the nascent VNC, are indicated in purple. (B, C) Late gastrula. Dorsal (B) and transverse sections (C). White outlines demarcate the NEC, and dashed orange outlines indicate the endomesoderm. (E, F) Early elongation. (E) Lateral maximum projection. White arrows indicate *Sce-hb9* expression in the trunk region. (F) Transverse section through the early elongation trunk highlighting the dorsoventral expression domains of *Sce-hb9* and *Sce-nk6* within the nascent VNC. White outlines demarcate the nascent VNC and orange outlines demarcate mesodermal cells and the intestine (dashed outline) (G, H) Hatchling. (G) Maximum projection shown in lateral (left) and dorsal (right) views. For anatomical context, refer to Fig. 2A’’. (H) Transverse section through the hatchling trunk highlighting the dorsoventral expression domains of *Sce-hb9* and *Sce-nk6* within the lateral somata clusters (purple outline). Orange outlines demarcate mesodermal derivatives, including the longitudinal muscle cells, and intestine (dashed outline) . The asterisk marks the position of the mouth. Scale bars: 50 µm, except (G): 100 µm. Orientation is indicated in the schematic representations and, where applicable, in the upper right corner of each panel. Abbreviations: ect, ectoderm; eme, endomesoderm; epi, epidermis; hb, head bud; in, intestine; lms, longitudinal muscle somata; mes, mesodermal cells; mvs, medioventral somata clusters; nbs, neural cells of the developing VNC; nec, neuroectoderm; sto, stomodeum; tb, tail bud.

### Expression of *th* and *dbh*

At hatching, *Sce-th* is expressed in a single bilateral pair of neurons in the anterior lateral somata clusters, with a second pair appearing in the early juveniles (Fig. 6A, B). *Sce-th* expression is also detected in the vestibular ganglia in the early juvenile (inset in Fig. 6B). *Sce-dbh* expression is not detected in hatchlings. *Sce-dbh* expression was not detected in hatchlings (data not shown). In early juveniles, *Sce-dbh* expression appears in three bilateral pairs of cells within the anterior lateral somata clusters and in cells of the anterior mouth (Fig. 6C). The vestibular ganglia also show a low-level *Sce-dbh* expression (inset in Fig. 6B).

**Figure 6.**
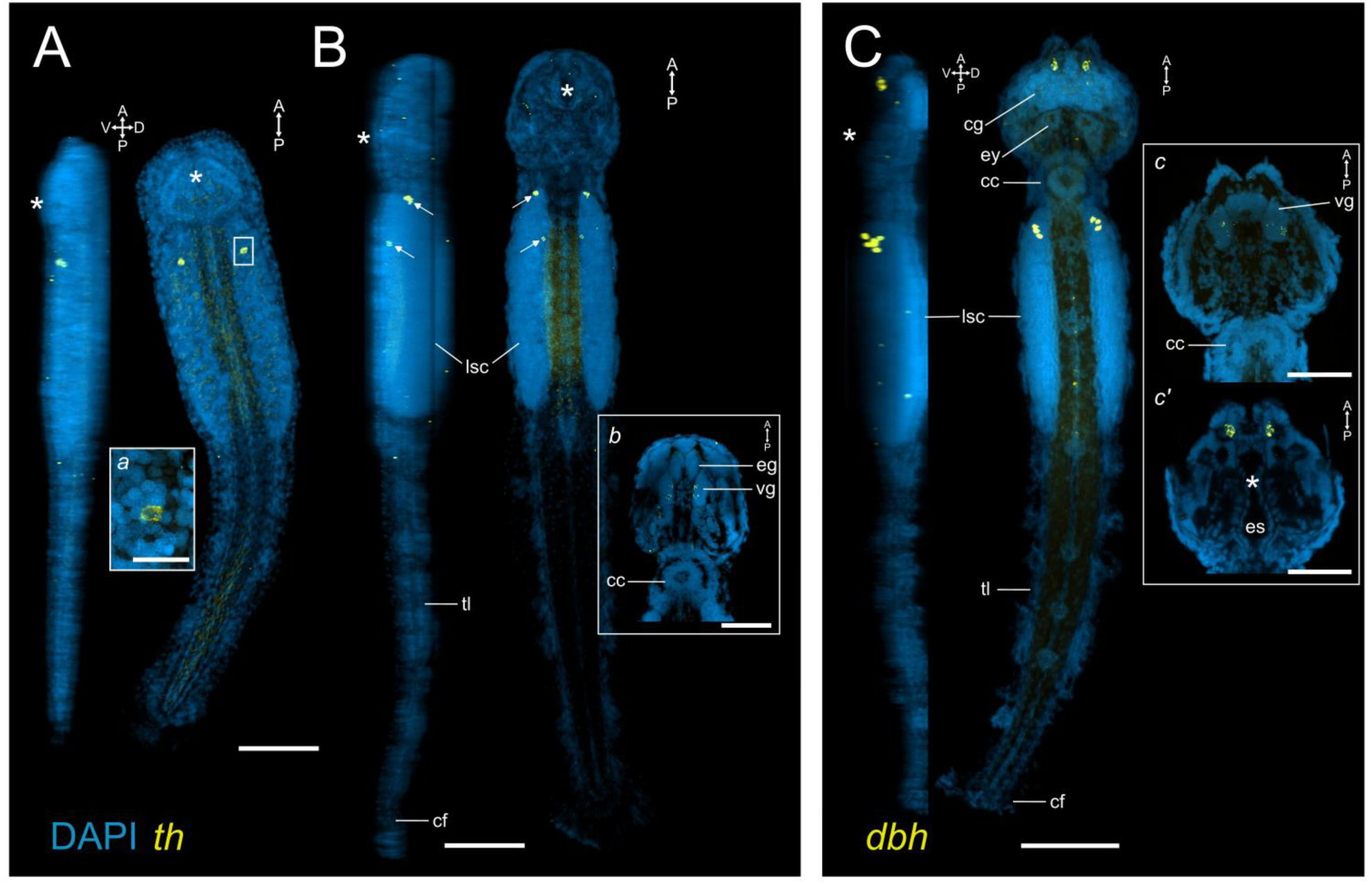
Expression patterns of *Sce-th* and *Sce-dbh* during early post-embryonic development of *Spadella cephaloptera* (A, B) *Sce-th* expression. (A) Maximum projection of a hatchling in lateral (left) and dorsal view (right). Inset *a* shows a magnified view of one of the two *Sce-th^+^* cells. For anatomical context, refer to Figure 2’’. (B) Maximum-intensity projection of an early juvenile in lateral (left) and dorsal view (right). White arrows indicate *Sce-th*⁺ cells within the lateral somata clusters. Inset *b* shows a dorsal section of the head highlighting *Sce-th* expression in the vestibular ganglia. The asterisk marks the position of the mouth. (C) *Sce-dbh* expression. Maximum projection of an early juvenile in lateral (left) and dorsal view (right). Insets *c* and *c′* show dorsal sections of the head at different dorsoventral levels. Inset *c* corresponds to a more dorsal section highlighting *Sce-dbh* expression in the vestibular ganglia, whereas inset *c′* shows a more ventral level encompassing the anterior mouth region. Scale bars: 100 µm, except (a): 25 µm; (b, c, c′): 50 µm. Orientation is indicated in the upper right corner of each panel. Abbreviations: cc, corona cilata; cf, caudal fin; cg, cerebral ganglion; eg, esophageal ganglia; ey, eye; es, esophagus; lsc, lateral somata clusters; tl, tail; vg,vestibular ganglia.

## Discussion

This study examines the spatial and temporal domains of broadly conserved developmental genes associated with nervous system formation in the chaetognath *Spadella cephaloptera*. Spiralian nervous systems span substantial architectural diversity, and chaetognaths possess a compact nervous system with a comparatively distinctive gross organization (Harzsch and Wanninger, 2010; Rieger et al., 2011, 2010), thereby providing a valuable context for assessing how conserved molecular patterning signatures are spatiotemporally deployed within this lineage.

Our analyses identify molecularly distinguishable domains corresponding to early neurogenic territories, dorsoventral patterning context, and the initial differentiation of neuronal subtypes (Fig. 7). These expression-based data map the emergence of discrete neural territories and neuronal subtypes in *S. cephaloptera*, highlighting both commonly observed bilaterian patterns and lineage-specific features of nervous system development.

**Figure 7.**
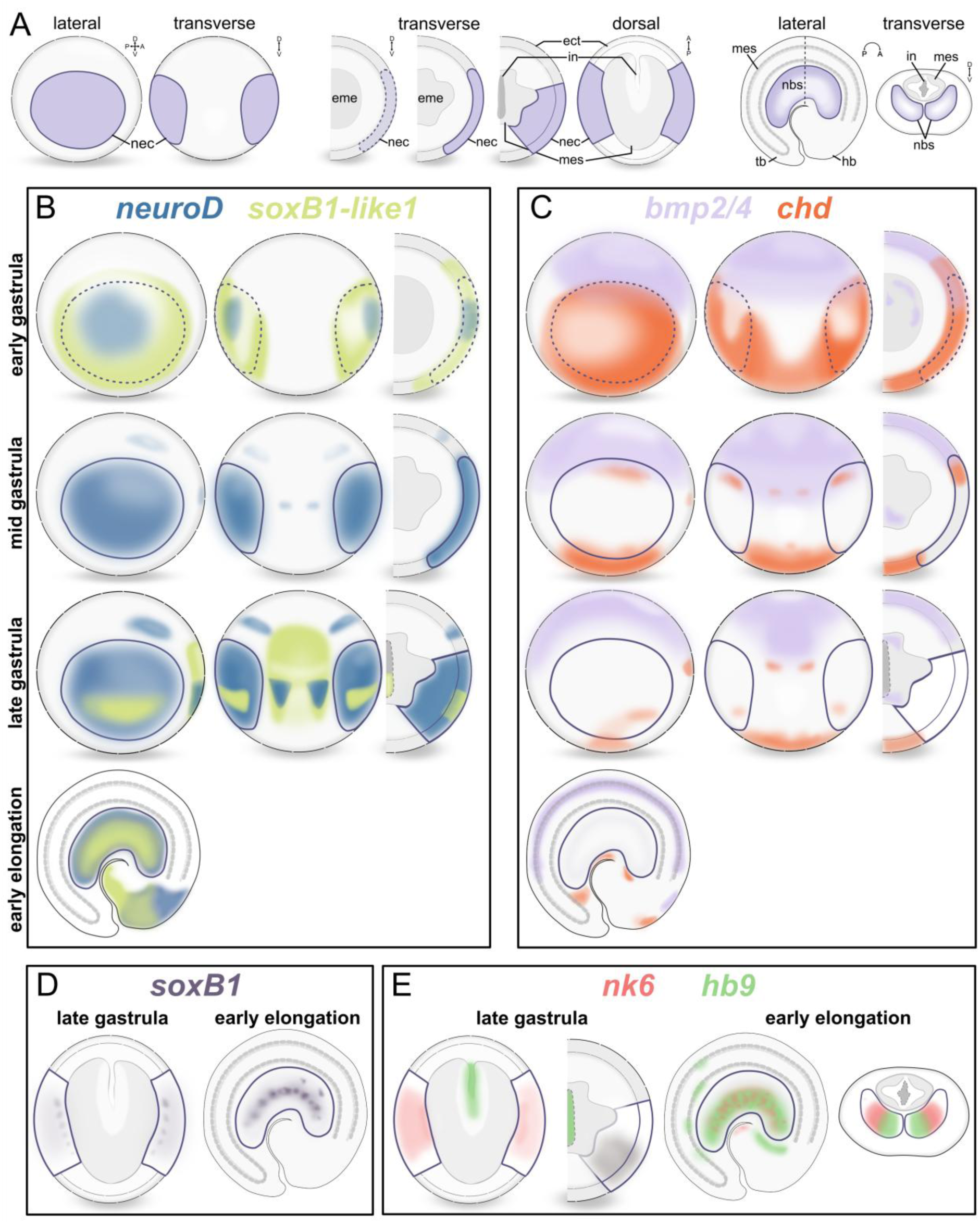
Schematic representation of neural-gene associated and dorsoventral patterning gene-expression domains during embryonic development of *Spadella cephaloptera*. (A) Anatomical reference schematics showing whole gastrula (lateral and dorsal views) and representative sections (transverse sections of early, mid, and late gastrula; dorsal section of late gastrula; and lateral and transverse sections through the trunk at early elongation). Neurogenic regions, including the neuroectoderm (NEC) and neural cells of the developing ventral nerve cord (VNC), are indicated in purple. At the early gastrula stage, the purple domain corresponds to the presumptive neuroectoderm. (B) Summary of *Sce-soxB1-like1* and *Sce-neuroD* expression domains across gastrula and early elongation stages. (C) Summary of *Sce-bmp2/4* and *Sce-chordin* expression domains across gastrula and early elongation stages. (D) Summary of *Sce-soxB1* expression domains in late gastrula and early elongation. (E) Summary of *Sce-nk6* and *Sce-hb9* expression domains in late gastrula and early elongation.

### Mapping early neurogenic territories in *Spadella*

Classical embryological descriptions of *Spadella cephaloptera* noted how the cells of the ventral nerve center originate from the ventrolateral ectoderm of the embryo, but did not resolve the identity of the neuroectodermal precursors that give rise to these neurons, nor clarify how early neurogenic cell populations are established (John, 1933). Here, we integrate *Sce-soxB1-like1* and *Sce-neuroD* expression with DAPI-based nuclear morphology to map the emergence and regionalization of the neuroectoderm during *S. cephaloptera* early development (Fig. 3; Fig. 7B).

The broad domain of *Sce-soxB1-like1* expression in the early gastrula spatially coincides with an ectodermal region characterized by comparatively large, relatively DAPI-faint nuclei and frequent mitotic figures observed in this population during gastrulation. Although reduced DAPI intensity and large nuclei may reflect a less condensed chromatin state and have been associated in some systems with progenitor or proliferative populations (Faro-Trindade and Cook, 2006; Madl et al., 2020), these features alone are not sufficient to establish progenitor identity. We therefore interpret this SoxB1-positive territory as a putative early neurogenic or neural-competent domain, based primarily on its spatial extent of SoxB1-class expression and anatomical context. Consistent with this interpretation, SoxB1 orthologs are widely implicated in neuroectodermal competence and early progenitor-like states across bilaterians (e.g., Sox1/2/3 in vertebrates and SoxNeuro in arthropods), including in diverse spiralian lineages (Buescher et al., 2002; Bylund et al., 2003; Deryckere et al., 2021; Hartenstein and Stollewerk, 2015; Kerner et al., 2009; Kurtova et al., 2024; Monjo and Romero, 2015b).

Notably, weak *Sce-neuroD* signal is already detectable within this same territory, reinforcing the view that this region represents an early neurogenic domain. This lateral domain is also positionally continuous with the *Sce-neuroD^+^* ventrolateral NEC regions observed in later stages.

### Early neuroectoderm proliferation and neural specification

The present study shows that by mid-gastrulation, the NEC contains dividing cells within the LNE population, and that SNE cells emerge, occurring adjacent to and interspersed among LNE cells. *Sce-neuroD* expression first appears broadly across the NEC at this stage, spanning both LNE and SNE populations, suggesting that a proneural-associated transcriptional program is active across the neuroectoderm. Because *Sce-soxB1-like1* expression was not characterized at mid-gastrula, it limits our ability to assign temporal relationships between broad *SoxB1-like1* expression and proneural-associated transcription at this stage.

By late gastrulation, the two markers show a more resolved spatial relationship, with *Sce-neuroD* expression maintained across the neuroectoderm and *Sce-soxB1-like1* restricted to the ventral portion of the NEC, where both genes co-localize within the LNE population (Fig. 3G – I; Fig. 7B). *Sce-neuroD* expression can also be detected in mitotically active neuroectodermal cells (Fig. 3F, I), supporting a neurogenic configuration in which *neuroD* expression initiates prior to cell-cycle exit and overlaps with mitotic activity in the neuroectoderm. Comparable patterns have been reported in several spiralian taxa, including the annelid *Platynereis dumerilii* and planarians, where *neuroD* orthologs are expressed within proliferative neuroectodermal or progenitor populations (Cowles et al., 2013; Monjo and Romero, 2015a; Simionato et al., 2008).

By contrast, in other bilaterians such as the annelid *Capitella teleta*, the cephalopod *Octopus vulgaris*, and vertebrates, NeuroD-family genes are predominantly associated with terminally dividing or post-mitotic neural precursors (Bertrand et al., 2002; Deryckere et al., 2021; Sur et al., 2017, 2020). This suggests that, although the activation timing of NeuroD-family genes within the neurogenic program varies among bilaterian lineages, early expression within proliferative neural territories is a recurring pattern in several taxa. The *Sce-neuroD* expression profile fits within this broader spiralian diversity, expanding the comparative evidence that multiple neurogenic modes involving early-onset *neuroD* activity occur across Spiralia.

SNE cells also increase in number and migrate inward (potentially through ingression) during late gastrulation, accumulating near the ventrolateral borders of the endomesoderm (Fig. 1H). This transition coincides with the emergence of detectable *Sce-soxB1* expression within a subset of the inner SNE population, arranged in a repeated pattern along the anterior–posterior axis rather than uniformly across the neuroectoderm (Fig. 2; Fig. 7D). A comparable patterning is maintained during early elongation within the developing lateral somata clusters, where *Sce-soxB1* marks a specific subpopulation of neural cells. This spatially restricted deployment contrasts with the broad progenitor-associated expression reported for some bilaterian SoxB1 homologs, raising the possibility that *Sce-soxB1* reflects lineage- or paralog-specific deployment. Consistent with this interpretation, SoxB1 paralogs in several bilaterians exhibit partitioned spatiotemporal expression while retaining broadly conserved roles in neural development. For example, in zebrafish, *sox1a* and *sox1b* show later-onset and regionally restricted expression relative to other SoxB1 paralogs (Okuda et al., 2006), while in amphioxus, *soxB1a* and *soxB1c* display distinct early expression profiles that later diverge across neural and non-neural tissues (Meulemans and Bronner-Fraser, 2007). A similar temporal partitioning has been reported in the planarian *Schmidtea polychroa*, where *soxB1-1* is activated earlier in presumptive neural progenitors and *soxB1-2* appears later during differentiation (Monjo and Romero, 2015a). Together, these comparisons provide context for interpreting the stage- and region-specific deployment of SoxB1 genes in *S. cephaloptera*.

In the early elongation stage, *Sce-soxB1-like1* expression becomes broadly distributed across the nascent VNC, extending beyond its earlier restriction to the LNE domain. This broad deployment is maintained into the hatchling stage, consistent with an expansion from its putative neural progenitor domain to a more widespread neural expression profile. While *Sce-neuroD* remains broadly expressed within the developing ventral nerve center, overlapping with the *Sce-Soxb1-like1*, it is also detected a discrete population of dorsal epidermal cells in the head bud, distinguishable by their comparatively DAPI signal (Fig. S10H). This anterior, extra-neural *Sce-neuroD* domain persists into the hatchling, where expression is observed in the eyes and the corona ciliata. Although the cellular identity and lineage relationships of the *Sce-neuroD⁺* cells cannot be resolved with our data, their persistence in anterior sensory-associated structures is consistent with a potential role for NeuroD in chaetognath sensory organ development.

### Molecular characterization of a putative anterior neuroectodermal territory

We identify a discrete molecular territory marked by *Sce-neuroD* and *Sce-soxB1-like1* during late gastrulation and early elongation. In the late gastrula, *Sce-neuroD* expression forms paired anterior domains that overlap with the broader anterior ectodermal expression of *Sce-soxB1-like1* (Fig. 3G – I; Fig. 7A). Notably, this territory is positioned within the anterior ectodermal field marked by *Sce-otx*, providing additional positional support for a potential anterior neuroectodermal identity (Ordoñez and Wollesen, 2025). With the onset of axial elongation, a bilateral *Sce-neuroD⁺*/*Sce-soxB1-like1*⁺ territory is observed on both sides of the anterior head bud (Fig. S10H). This position closely matches the thickened anterior neuroectoderm in the early elongation stage described in classical embryological studies as contributing to the cerebral ganglia (John, 1933). However, definitive assignment of this territory to the presumptive head neuroectoderm will require experimental validation (e.g.) lineage-tracing.

### Overlapping proliferative and differentiation-associated transcriptional signatures in the hatchling nervous system

Ultrastructural and immunohistochemical context reveals that *S. cephaloptera* hatchlings emerge with a nervous system that is functionally competent yet developmentally immature. Rieger et al. (2011) described the VNC containing differentiated neurons, including RFamide+ and synapsin+ populations, while noting that synaptic networks, commissural tracts, and vesicle complements remain sparse and continue to expand post-hatching. Our molecular data are consistent with this interpretation, as the same neural regions express transcriptional markers spanning multiple neurogenic states, including early neurogenic programs (*Sce-neuroD*, *Sce-soxB2*) and markers associated with neuronal maturation (*Sce-hb9*, *Sce-th*, *Sce-dbh*). Broad *Sce-elav* expression in the hatchling VNC and brain (Ordoñez and Wollesen, 2024) further supports the presence of neurons that have entered molecular stages of differentiation.

In addition, the hatchling brain and VNC remain mitotically active for several days post-hatching, with BrdU⁺ nuclei forming bilateral bands along the inner margins of the lateral somata clusters (Perez et al., 2013). When considered alongside our molecular data, these proliferative cells correspond to neural populations expressing gene complements typically associated with early neurogenic or fate-biased precursor states rather than naïve neuroectoderm. This configuration contrasts with many metazoan systems, including cnidarians, insects, chordates, and *Capitella*, in which NeuroD- and Elav-family genes are largely restricted to post-mitotic differentiating neurons (D’Amico et al., 2013; Kim et al., 1996; Nakanishi et al., 2012; Robinow and White, 1991; Sur et al., 2017). The coexistence of proliferative activity and differentiation-associated markers is therefore consistent with a population of transit-amplifying or lineage-restricted neural precursors interspersed with differentiating neurons, rather than a large undifferentiated progenitor pool. Whether these proliferative cells derive from persisting embryonic neuroectoderm or represent a distinct progenitor population remains unresolved. Overall, the hatchling VNC integrates proliferative, differentiating, and mature neuronal features within a compact neural architecture that continues to develop after hatching.

In this context, the differential deployment of three SoxB paralogs provides molecular insight into the neurogenic landscape of the hatchling CNS. *Sce-soxB1-like1* is broadly expressed across hatchling neural structures, consistent with activity in differentiating neural territories. This pattern aligns with observations from multiple bilaterian systems in which SoxB1-type genes function not only in neural competence but also during later stages of neural differentiation (Buescher et al., 2002; Deryckere et al., 2021; Ekonomou et al., 2005; Ferrero et al., 2014; Monjo and Romero, 2015a; Overton et al., 2002; Sur et al., 2017; Vidal et al., 2015). By contrast, *Sce-soxB1* shows a more spatially restricted expression pattern, being localized to the presumptive anteroventral cephalic ganglion anlage and to discrete subpopulations of the lateral somata clusters, with weaker expression in parts of the cerebral ganglion and medioventral somata clusters. This distribution suggests that *Sce-soxB1* is enriched in specific neural subdomains rather than broadly marking differentiated neurons, potentially reflecting roles in neural lineage refinement or regional specification. Unlike the two SoxB1 paralogs, *Sce-soxB2* expression is first detected at the hatchling stage, specifically within the lateral somata clusters and the developing ventral cephalic ganglia. This late onset is consistent with the deployment of SoxB2 homologs in several bilaterian lineages, where these transcription factors are more frequently associated with neural differentiation and subtype specification rather than progenitor maintenance (Guth and Wegner, 2008; Monjo and Romero, 2015a; Overton et al., 2002; Wegner and Stolt, 2005; Zhao and Skeath, 2002). Thus, the distinct expression profiles of *Sce-soxB1-like1*, *Sce-soxB1*, and *Sce-soxB2* across neural territories exhibiting features of both proliferative activity and maturation provide pattern-level support for the idea that SoxB family members can act at multiple stages of neurogenesis, as proposed in other bilaterian systems.

### Bmp2/4-chordin dorsoventral patterning during early neuroectodermal specification

In many bilaterians, BMP signaling, modulated by antagonists of the Chordin family and related BMP inhibitors, constitutes a core dorsoventral (DV) patterning system and is often associated with the positioning of neuroectodermal territories relative to the surrounding ectoderm (Bier and De Robertis, 2015; Erwin, 2009). In *S. cephaloptera, Sce-bmp2/4* and *Sce-chordin* exhibit a reciprocal DV expression pattern in the early gastrula, coinciding with the stage at which the neuroectodermal markers, *Sce-soxB1-like1* and *Sce-neuroD,* are first detected in our dataset (Fig. 4; Fig. 7C). These neural markers arise within the lateral to ventrolateral regions of the embryo that correspond to *Sce-chordin*^+^ and *Sce-bmp2/4*-reduced domains. The BMP pathway activity at this stage is reflected by dorsally localized phosphorylated Smad1/5/8 (pSmad1/5/8), indicating active BMP signaling output (Knabl et al., 2025). Notably, pSmad1/5/8 does not fully coincide with *Sce-bmp2/4* transcript domains, a pattern that is expected in BMP–Chordin systems, where extracellular modulation, including ligand diffusion and antagonistic interactions, shape signaling fields that extend beyond sites of ligand (i.e., *Sce-bmp2/4*) transcription (Rahman et al., 2015; Zakin and De Robertis, 2010). These observations indicate the presence of a dorsoventrally polarized BMP signaling environment during gastrulation, at the time early neurogenic territories become molecularly distinguishable.

This arrangement parallels patterns in several bilaterian systems, including lophotrochozoans, insects, and vertebrates, in which regions of relatively low BMP activity or elevated *chordin* expression are frequently associated with territories permissive for neural competence, whereas BMP-high domains typically correspond predominantly to non-neural ectoderm (Carrillo-Baltodano et al., 2025; Denes et al., 2007; Holley et al., 1995; Lewin et al., 2024; Martín-Durán et al., 2018; Tan et al., 2022). At the same time, substantial variability of DV patterning mechanisms has also been reported across spiralian lineages, including cases in which canonical BMP–Chordin antagonism is modified or replaced (Kuo and Weisblat, 2011; Lanza and Seaver, 2020, 2018; Lyons et al., 2020).

As gastrulation proceeds, DV expression domains in *S. cephaloptera* become narrower, with *Sce-chordin* expression progressively retracting ventrally and *Sce-bmp2/4* persisting dorsally and in endomesodermal cells. By early elongation, these genes no longer form broad axial gradients and, by the hatchling stage, both genes are restricted to discrete cell populations. This progression indicates that DV patterning in *S. cephaloptera* is temporally restricted to gastrulation, coinciding with the establishment of initial neurogenic territories, and that *Sce-bmp2/4* and *Sce-chordin* are subsequently redeployed in more localized and tissue-specific roles.

Importantly, BMP signaling is not universally excluded from neural territories and can be active within developing nervous systems of cnidarians and bilaterians, including *S. cephaloptera* (Knabl et al., 2025; Moore et al., 2025; Mörsdorf et al., 2024; Pasini et al., 2006; Rusten et al., 2002; Waki et al., 2015). In several systems, BMP activity has been implicated in aspects of neurogenic regulation, including neuronal differentiation or subtype patterning, rather than acting solely as an anti-neural signal (Denes et al., 2007; Knabl et al., 2025; Lambert et al., 2016; McClay et al., 2018; Tan et al., 2022). Taken together, our data support a model in which *S. cephaloptera* shows an early and transient DV BMP–Chordin expression framework associated with neuroectodermal emergence, followed by a later, more localized deployment of *Sce-chordin*, *Sce-bmp2/4*, and associated BMP pathway components within specific tissues, including neural territories.

### *Nk6* and *hb9* expression patterns indicate early motor neuron-associated ventral patterning

To investigate how ventral neuronal subtypes emerge in *Spadella cephaloptera*, we characterized the expression patterns of *Sce-nk6* and *Sce-hb9*, two conserved transcription factors widely implicated in ventral CNS patterning and motor neuron specification across bilaterians (Arber et al., 1999; Cheesman et al., 2004; Denes et al., 2007; Dichmann and Harland, 2011; Ferrier et al., 2001; Odden et al., 2002; Schmidt et al., 1997). In *S. cephaloptera*, *Sce-nk6* expression is detectable by late gastrulation in a subset of SNE cells within the midventral neuroectoderm, indicating that molecular subdivision within the neurogenic field is already initiated at this stage (Fig. 5; Fig. 7D). This ventral bias becomes more apparent during early elongation, when *Sce-nk6* and *Sce-hb9* jointly mark the developing VNC in partially overlapping domains. In hatchlings, *Sce-hb9* expression is detected in the presumptive anteroventral cephalic ganglion anlage, consistent with previous interpretations that the vestibular ganglia accommodate motor neurons innervating the head musculature of chaetognaths (Müller et al., 2018; Rieger et al., 2011; Shinn, 1997). Notably, within the lateral somata clusters, a predominantly ventral *Sce-hb9* domain occupies a subset of a broader Sce*-nk6^+^*territory. This overlapping *Sce-nk6* ⁺/ *Sce-hb9*⁺ organization is consistent with conserved ventral CNS patterning schemes, in which Nk6-family transcription factors mark ventral progenitor or precursor domains that give rise to motor neurons and specific ventral interneuron subtypes, while *hb9*/*mnx1* (*motor neuron and pancreas homeobox 1*) functions as a terminal selector for motor neuron identity (Arber et al., 1999; Cheesman et al., 2004; Denes et al., 2007; Dichmann and Harland, 2011; Ferrier et al., 2001; Odden et al., 2002; Schmidt et al., 1997). Moreover, the spatial deployment of *Sce-nk6* and *Sce-hb9* during development further provides molecular support for a mediolateral organization of the VNC previously proposed for *S. cephaloptera* (Ordoñez and Wollesen, 2024).

Although our analysis focuses on neural territories, *Sce-hb9* is also expressed in the presumptive endoderm at late gastrulation and later in the hatchling gut. Comparable dual expression of *hb9*/*mnx1* expression to both the CNS and the gut has been reported in *Drosophila* (Odden et al., 2002) and across deuterostomes, including sea urchin, amphioxus, and vertebrates (Bellomonte et al., 1998; Harrison et al., 1994, 1999). This shared neural–endodermal expression pattern therefore appears to represent an evolutionarily conserved feature of *hb9/mnx1* deployment across Bilateria (Bellomonte et al., 1998; Ferrier et al., 2001; Harrison et al., 1999, 1994; Saha et al., 1997).

### Post-embryonic deployment of catecholamine pathway components

Tyrosine hydroxylase (TH) and dopamine β-hydroxylase (DBH) encode core enzymes in catecholamine biosynthesis, and their biochemical activities are conserved throughout Bilateria (Goulty et al., 2023; Sloley and Juorio, 1995). Both enzymes have been widely used as molecular markers associated with monoaminergic neuron identity, including dopaminergic and noradrenergic neurons, in well-characterized systems (Flames and Hobert, 2011). In the *S. cephaloptera* hatchling, *Sce-th* expression is detected in a small bilateral subset of the VNC cells, whereas *Sce-dbh* is not detected at this stage. *Sce-dbh* expression is first detected in early juveniles, in the anterior VNC and head region. This temporal offset suggests that catecholamine-associated molecular components are activated sequentially during early post-embryonic development. In the early juvenile, *Sce-dbh* is restricted to the anterior-most region of the VNC with additional non-neural expression in the head region. While this region may later contribute to anterior mouth structures and associated sensory elements, the cellular identity and functional significance of these anterior head domains remain unresolved at this stage.

At present, direct comparisons with other spiralians remain limited due to the scarcity of molecular-level data on DBH deployment in this group. Non-neuronal *dbh* expression has only been reported in a small number of spiralian species, including hemocytes and liver tissue in bivalve mollusks (Li et al., 2021; Zhou et al., 2011) and protein-level localization in the urn cells of the dicyemid *Dicyema japonicum* (Lu et al., 2019). Nevertheless, these observations place the *S. cephaloptera* data within a broader but still sparse comparative context and support the conclusion that *Sce-th* and *Sce-dbh* show temporally and spatially distinct expression patterns during early post-embryonic development.

## Conclusion

Our analysis shows that nervous system development in *Spadella cephaloptera* relies on conserved bilaterian molecular programs that are deployed in lineage-specific spatial and temporal patterns. *Sce-soxB* and *Sce-neuroD* expression delineates a neuroectodermal territory in which proliferative and differentiation-associated programs overlap, while a transient *Sce-bmp2/4* - *Sce-chordin* dorsoventral pattern coincides with the emergence of early neurogenic territories. Ventral neuronal subtype patterning, including early motor-neuron specification marked by *Sce-nk6* and *Sce*-*hb9*, further supports the conservation of core bilaterian neural patterning mechanisms in chaetognaths. In contrast, the restricted expression of *Sce-th* and the delayed onset of *Sce-dbh* indicate that catecholaminergic pathway components are assembled progressively within established neural territories rather than as a unified monoaminergic system. Collectively, these findings define the spatial and temporal organization of early neurogenesis in chaetognaths and provide a comparative reference for future analyses of how markers associated with bilaterian neurogenesis are deployed across spiralian lineages.

## Declarations

### Data availability statement

The assembled transcriptome for *Spadella cephaloptera* is accessible on Zenodo (https://zenodo.org/record/7602960#.Y90U0oSZOUk/DOI:10.5281/zenodo.7602960). Newly obtained *S. cephaloptera* sequences used in the phylogenetic analyses have been deposited to GenBank (accession numbers: PZ054574 - PZ054584).

### Authors’ contributions

J.F.O. designed and performed the experiments, maintained and fixed samples, performed the analyses, and wrote the manuscript. A.F. and C.C.G.B. collected animals, maintained and fixed samples, reviewed and edited the manuscript. T.W. designed the experiments, secured funding, supervised, and contributed to analyzing the results and to writing of the manuscript. All authors have read and approved of the final version of the manuscript.

## Acknowledgements

The authors are grateful for financial support by the Faculty of Life Sciences of the University of Vienna.

## References

1. Altschul, S.F., Gish, W., Miller, W., Myers, E.W., Lipman, D.J., 1990. Basic local alignment search tool. J. Mol. Biol. 215, 403–410. 10.1016/S0022-2836(05)80360-2

2. Arber, S., Han, B., Mendelsohn, M., Smith, M., Jessell, T.M., Sockanathan, S., 1999. Requirement for the Homeobox Gene Hb9 in the Consolidation of Motor Neuron Identity. Neuron 23, 659–674. 10.1016/S0896-6273(01)80026-X

3. Arendt, D., Denes, A.S., Jékely, G., Tessmar-Raible, K., 2008. The evolution of nervous system centralization. Philos. Trans. R. Soc. B Biol. Sci. 363, 1523–1528. 10.1098/rstb.2007.2242

4. Arendt, D., Tosches, M.A., Marlow, H., 2016. From nerve net to nerve ring, nerve cord and brain — evolution of the nervous system. Nat. Rev. Neurosci. 17, 61–72. 10.1038/nrn.2015.15

5. Bellomonte, D., Di Bernardo, M., Russo, R., Caronia, G., Spinelli, G., 1998. Highly restricted expression at the ectoderm–endoderm boundary of PIHbox 9, a sea urchin homeobox gene related to the human HB9 gene. Mech. Dev. 74, 185–188. 10.1016/S0925-4773(98)00064-1

6. Bertrand, N., Castro, D.S., Guillemot, F., 2002. Proneural genes and the specification of neural cell types. Nat. Rev. Neurosci. 3, 517–530. 10.1038/nrn874

7. Bier, E., De Robertis, E.M., 2015. BMP gradients: A paradigm for morphogen-mediated developmental patterning. Science 348, aaa5838. 10.1126/science.aaa5838

8. Bruce, H.S., Jerz, G., Kelly, S.R., McCarthy, J., Pomerantz, A., Senevirathne, G., Sherrard, A., Sun, D.A., Wolff, C., Patel, N., 2021. Hybridization Chain Reaction (HCR) In Situ Protocol v1. 10.17504/protocols.io.bunznvf6

9. Buescher, M., Hing, F.S., Chia, W., 2002. Formation of neuroblasts in the embryonic central nervous system of *Drosophila melanogaster* is controlled by *SoxNeuro*. Development 129, 4193–4203. 10.1242/dev.129.18.4193

10. Buresi, A., Andouche, A., Navet, S., Bassaglia, Y., Bonnaud-Ponticelli, L., Baratte, S., 2016. Nervous system development in cephalopods: How egg yolk-richness modifies the topology of the mediolateral patterning system. Dev. Biol. 415, 143–156. 10.1016/j.ydbio.2016.04.027

11. Bylund, M., Andersson, E., Novitch, B.G., Muhr, J., 2003. Vertebrate neurogenesis is counteracted by Sox1–3 activity. Nat. Neurosci. 6, 1162–1168. 10.1038/nn1131

12. Carrillo-Baltodano, A.M., Haillot, E., Meha, S.M., Luqman, I., Pashaj, A., Lee, Y.-J., Lu, T.-M., Ferrier, D.E.K., Schneider, S.Q., Martín-Durán, J.M., 2025. Developmental system drift in dorsoventral patterning is linked to transitions to autonomous development in Annelida. 10.1101/2025.05.29.656861

13. Cheesman, S.E., Layden, M.J., Von Ohlen, T., Doe, C.Q., Eisen, J.S., 2004. Zebrafish and fly Nkx6 proteins have similar CNS expression patterns and regulate motoneuron formation. Development 131, 5221–5232. 10.1242/dev.01397

14. Choi, H.M.T., Beck, V.A., Pierce, N.A., 2014. Next-Generation *in Situ* Hybridization Chain Reaction: Higher Gain, Lower Cost, Greater Durability. ACS Nano 8, 4284–4294. 10.1021/nn405717p

15. Choi, H.M.T., Schwarzkopf, M., Fornace, M.E., Acharya, A., Artavanis, G., Stegmaier, J., Cunha, A., Pierce, N.A., 2018. Third-generation in situ hybridization chain reaction: multiplexed, quantitative, sensitive, versatile, robust. 10.1101/285213

16. Cowles, M.W., Brown, D.D.R., Nisperos, S.V., Stanley, B.N., Pearson, B.J., Zayas, R.M., 2013. Genome-wide analysis of the bHLH gene family in planarians identifies factors required for adult neurogenesis and neuronal regeneration. Development 140, 4691–4702. 10.1242/dev.098616

17. D’Amico, L.A., Boujard, D., Coumailleau, P., 2013. The Neurogenic Factor NeuroD1 Is Expressed in Post-Mitotic Cells during Juvenile and Adult Xenopus Neurogenesis and Not in Progenitor or Radial Glial Cells. PLoS ONE 8, e66487. 10.1371/journal.pone.0066487

18. De Robertis, E.M., Sasai, Y., 1996. A common plan for dorsoventral patterning in Bilateria. Nature 380, 37–40. 10.1038/380037a0

19. Denes, A.S., Jékely, G., Steinmetz, P.R.H., Raible, F., Snyman, H., Prud’homme, B., Ferrier, D.E.K., Balavoine, G., Arendt, D., 2007. Molecular Architecture of Annelid Nerve Cord Supports Common Origin of Nervous System Centralization in Bilateria. Cell 129, 277–288. 10.1016/j.cell.2007.02.040

20. Deryckere, A., Styfhals, R., Elagoz, A.M., Maes, G.E., Seuntjens, E., 2021. Identification of neural progenitor cells and their progeny reveals long distance migration in the developing octopus brain. eLife 10, e69161. 10.7554/eLife.69161

21. Dichmann, D.S., Harland, R.M., 2011. Nkx6 genes pattern the frog neural plate and Nkx6.1 is necessary for motoneuron axon projection. Dev. Biol. 349, 378–386. 10.1016/j.ydbio.2010.10.030

22. Doncaster, L., 1902. On the Development of Sagitta; with Notes on the Anatomy of the Adult. J. Cell Sci. S2-46, 351–395. 10.1242/jcs.s2-46.182.351

23. Ekonomou, A., Kazanis, I., Malas, S., Wood, H., Alifragis, P., Denaxa, M., Karagogeos, D., Constanti, A., Lovell-Badge, R., Episkopou, V., 2005. Neuronal Migration and Ventral Subtype Identity in the Telencephalon Depend on SOX1. PLoS Biol. 3, e186. 10.1371/journal.pbio.0030186

24. Erwin, D.H., 2009. Early origin of the bilaterian developmental toolkit. Philos. Trans. R. Soc. B Biol. Sci. 364, 2253–2261. 10.1098/rstb.2009.0038

25. Faltine-Gonzalez, D., Havrilak, J., Layden, M.J., 2023. The brain regulatory program predates central nervous system evolution. Sci. Rep. 13, 8626. 10.1038/s41598-023-35721-4

26. Faro-Trindade, I., Cook, P.R., 2006. A Conserved Organization of Transcription during Embryonic Stem Cell Differentiation and in Cells with High C Value. Mol. Biol. Cell 17, 2910–2920. 10.1091/mbc.e05-11-1024

27. Ferrero, E., Fischer, B., Russell, S., 2014. SoxNeuro orchestrates central nervous system specification and differentiation in Drosophila and is only partially redundant with Dichaete. Genome Biol. 15, R74. 10.1186/gb-2014-15-5-r74

28. Ferrier, D.E.K., Brooke, N.M., Panopoulou, G., Holland, P.W.H., 2001. The Mnx homeobox gene class defined by HB9, MNR2 and amphioxus AmphiMnx. Dev. Genes Evol. 211, 103–107. 10.1007/s004270000124

29. Flames, N., Hobert, O., 2011. Transcriptional Control of the Terminal Fate of Monoaminergic Neurons. Annu. Rev. Neurosci. 34, 153–184. 10.1146/annurev-neuro-061010-113824

30. Goto, T., Katayama-Kumoi, Y., Tohyama, M., Yoshida, M., 1992. Distribution and development of the serotonin-and RFamide-like immunoreactive neurons in the arrowworm, Paraspadella gotoi (Chaetognatha). Cell Tissue Res. 267, 215–222. 10.1007/BF00302958

31. Goto, T., Yoshida, M., 1987. Nervous System in Chaetognatha, in: Ali, M.A. (Ed.), Nervous Systems in Invertebrates. Springer US, Boston, MA, pp. 461–481. 10.1007/978-1-4613-1955-9_16

32. Goulty, M., Botton-Amiot, G., Rosato, E., Sprecher, S.G., Feuda, R., 2023. The monoaminergic system is a bilaterian innovation. Nat. Commun. 14, 3284. 10.1038/s41467-023-39030-2

33. Guth, S.I.E., Wegner, M., 2008. Having it both ways: Sox protein function between conservation and innovation. Cell. Mol. Life Sci. 65, 3000–3018. 10.1007/s00018-008-8138-7

34. Harrison, K.A., Druey, K.M., Deguchi, Y., Tuscano, J.M., Kehrl, J.H., 1994. A novel human homeobox gene distantly related to proboscipedia is expressed in lymphoid and pancreatic tissues. J. Biol. Chem. 269, 19968–19975. 10.1016/S0021-9258(17)32115-4

35. Harrison, K.A., Thaler, J., Pfaff, S.L., Gu, H., Kehrl, J.H., 1999. Pancreas dorsal lobe agenesis and abnormal islets of Langerhans in Hlxb9-deficient mice. Nat. Genet. 23, 71–75. 10.1038/12674

36. Hartenstein, V., Stollewerk, A., 2015. The Evolution of Early Neurogenesis. Dev. Cell 32, 390–407. 10.1016/j.devcel.2015.02.004

37. Harzsch, S., Müller, C.H., 2007. A new look at the ventral nerve centre of Sagitta: implications for the phylogenetic position of Chaetognatha (arrow worms) and the evolution of the bilaterian nervous system. Front. Zool. 4, 14. 10.1186/1742-9994-4-14

38. Harzsch, S., Müller, C.H.G., Rieger, V., Perez, Y., Sintoni, S., Sardet, C., Hansson, B., 2009. Fine structure of the ventral nerve centre and interspecific identification of individual neurons in the enigmatic Chaetognatha. Zoomorphology 128, 53–73. 10.1007/s00435-008-0074-4

39. Harzsch, S., Wanninger, A., 2010. Evolution of invertebrate nervous systems: the Chaetognatha as a case study. Acta Zool. 91, 35–43. 10.1111/j.1463-6395.2009.00423.x

40. Hejnol, A., Lowe, C.J., 2015. Embracing the comparative approach: how robust phylogenies and broader developmental sampling impacts the understanding of nervous system evolution. Philos. Trans. R. Soc. Lond. B. Biol. Sci. 370, 20150045. 10.1098/rstb.2015.0045

41. Hoang, D.T., Chernomor, O., Von Haeseler, A., Minh, B.Q., Vinh, L.S., 2018. UFBoot2: Improving the Ultrafast Bootstrap Approximation. Mol. Biol. Evol. 35, 518–522. 10.1093/molbev/msx281

42. Holland, N.D., 2003. Early central nervous system evolution: an era of skin brains? Nat. Rev. Neurosci. 4, 617–627. 10.1038/nrn1175

43. Holley, S.A., Jackson, P.D., Sasai, Y., Lu, B., Robertis, E.M.D., Hoffmann, F.M., Ferguson, E.L., 1995. A conserved system for dorsal-ventral patterning in insects and vertebrates involving sog and chordin. Nature 376, 249–253. 10.1038/376249a0

44. Janssen, R., Andersson, E., Betnér, E., Bijl, S., Fowler, W., Höök, L., Leyhr, J., Mannelqvist, A., Panara, V., Smith, K., Tiemann, S., 2018. Embryonic expression patterns and phylogenetic analysis of panarthropod sox genes: insight into nervous system development, segmentation and gonadogenesis. BMC Evol. Biol. 18, 88. 10.1186/s12862-018-1196-z

45. John, C.C., 1933. Habits, Structure, and Development of Spadella Cephaloptera. J. Cell Sci. S2-75, 625–696. 10.1242/jcs.s2-75.300.625

46. Kalyaanamoorthy, S., Minh, B.Q., Wong, T.K.F., Von Haeseler, A., Jermiin, L.S., 2017. ModelFinder: fast model selection for accurate phylogenetic estimates. Nat. Methods 14, 587–589. 10.1038/nmeth.4285

47. Katoh, K., 2005. MAFFT version 5: improvement in accuracy of multiple sequence alignment. Nucleic Acids Res. 33, 511–518. 10.1093/nar/gki198

48. Katoh, K., 2002. MAFFT: a novel method for rapid multiple sequence alignment based on fast Fourier transform. Nucleic Acids Res. 30, 3059–3066. 10.1093/nar/gkf436

49. Kelava, I., Rentzsch, F., Technau, U., 2015. Evolution of eumetazoan nervous systems: insights from cnidarians. Philos. Trans. R. Soc. B Biol. Sci. 370, 20150065. 10.1098/rstb.2015.0065

50. Kerner, P., Simionato, E., Le Gouar, M., Vervoort, M., 2009. Orthologs of key vertebrate neural genes are expressed during neurogenesis in the annelid *Platynereis dumerilii*. Evol. Dev. 11, 513–524. 10.1111/j.1525-142X.2009.00359.x

51. Kim, C.-H., Ueshima, E., Muraoka, O., Tanaka, H., Yeo, S.-Y., Huh, T.-L., Miki, N., 1996. Zebrafish elav/HuC homologue as a very early neuronal marker. Neurosci. Lett. 216, 109–112. 10.1016/0304-3940(96)13021-4

52. Knabl, P., Ordoñez, J.F., Montenegro Cabrera, J.D., Wollesen, T., Genikhovich, G., 2025. The anti-neural role of BMP signaling is a side effect of its global function in dorsoventral patterning. 10.1101/2025.06.08.658475

53. Kuehn, E., Clausen, D.S., Null, R.W., Metzger, B.M., Willis, A.D., Özpolat, B.D., 2022. Segment number threshold determines juvenile onset of germline cluster expansion in *Platynereis dumerilii*. J. Exp. Zoolog. B Mol. Dev. Evol. 338, 225–240. 10.1002/jez.b.23100

54. Kuo, D.-H., Weisblat, D.A., 2011. A New Molecular Logic for BMP-Mediated Dorsoventral Patterning in the Leech Helobdella. Curr. Biol. 21, 1282–1288. 10.1016/j.cub.2011.06.024

55. Kurtova, A.I., Finoshin, A.D., Aparina, M.S., Gazizova, G.R., Kozlova, O.S., Voronova, S.N., Shagimardanova, E.I., Ivashkin, E.G., Voronezhskaya, E.E., 2024. Expanded expression of pro-neurogenic factor SoxB1 during larval development of gastropod Lymnaea stagnalis suggests preadaptation to prolonged neurogenesis in Mollusca. Front. Neurosci. 18, 1346610. 10.3389/fnins.2024.1346610

56. Lambert, J.D., Johnson, A.B., Hudson, C.N., Chan, A., 2016. Dpp/BMP2-4 Mediates Signaling from the D-Quadrant Organizer in a Spiralian Embryo. Curr. Biol. 26, 2003–2010. 10.1016/j.cub.2016.05.059

57. Lanza, A.R., Seaver, E.C., 2020. Functional evidence that Activin/Nodal signaling is required for establishing the dorsal-ventral axis in the annelid *Capitella teleta*. Development 147, dev189373. 10.1242/dev.189373

58. Lanza, A.R., Seaver, E.C., 2018. An organizing role for the TGF-β signaling pathway in axes formation of the annelid Capitella teleta. Dev. Biol. 435, 26–40. 10.1016/j.ydbio.2018.01.004

59. Laumer, C.E., Fernández, R., Lemer, S., Combosch, D., Kocot, K.M., Riesgo, A., Andrade, S.C.S., Sterrer, W., Sørensen, M.V., Giribet, G., 2019. Revisiting metazoan phylogeny with genomic sampling of all phyla. Proc. R. Soc. B Biol. Sci. 286, 20190831. 10.1098/rspb.2019.0831

60. Lewin, T.D., Shimizu, K., Liao, I.J.-Y., Chen, M.-E., Endo, K., Satoh, N., Holland, P.W.H., Wong, Y.H., Luo, Y.-J., 2024. Brachiopod genome unveils the evolution of the BMP–Chordin network in bilaterian body patterning. 10.1101/2024.05.28.596352

61. Li, Z., Peng, M., Power, D.M., Niu, D., Dong, Z., Li, J., 2021. RNAi-mediated knock-down of the dopamine beta-hydroxylase gene changes growth of razor clams. Comp. Biochem. Physiol. B Biochem. Mol. Biol. 252, 110534. 10.1016/j.cbpb.2020.110534

62. Lowe, C.J., Wu, M., Salic, A., Evans, L., Lander, E., Stange-Thomann, N., Gruber, C.E., Gerhart, J., Kirschner, M., 2003. Anteroposterior Patterning in Hemichordates and the Origins of the Chordate Nervous System. Cell 113, 853–865. 10.1016/S0092-8674(03)00469-0

63. Lu, T.-M., Furuya, H., Satoh, N., 2019. Gene expression profiles of dicyemid life-cycle stages may explain how dispersing larvae locate new hosts. Zool. Lett. 5, 32. 10.1186/s40851-019-0146-y

64. Lyons, D.C., Perry, K.J., Batzel, G., Henry, J.Q., 2020. BMP signaling plays a role in anterior-neural/head development, but not organizer activity, in the gastropod Crepidula fornicata. Dev. Biol. 463, 135–157. 10.1016/j.ydbio.2020.04.008

65. Madl, C.M., LeSavage, B.L., Khariton, M., Heilshorn, S.C., 2020. Neural Progenitor Cells Alter Chromatin Organization and Neurotrophin Expression in Response to 3D Matrix Degradability. Adv. Healthc. Mater. 9, 2000754. 10.1002/adhm.202000754

66. Marlétaz, F., Peijnenburg, K.T.C.A., Goto, T., Satoh, N., Rokhsar, D.S., 2019. A New Spiralian Phylogeny Places the Enigmatic Arrow Worms among Gnathiferans. Curr. Biol. 29, 312–318.e3. 10.1016/j.cub.2018.11.042

67. Martín-Durán, J.M., Pang, K., Børve, A., Lê, H.S., Furu, A., Cannon, J.T., Jondelius, U., Hejnol, A., 2018. Convergent evolution of bilaterian nerve cords. Nature 553, 45–50. 10.1038/nature25030

68. Martinez, P., Bailly, X., Sprecher, S.G., Hartenstein, V., 2024. The Acoel nervous system: morphology and development. Neural Develop. 19, 9. 10.1186/s13064-024-00187-1

69. McClay, D.R., Miranda, E., Feinberg, S.L., 2018. Neurogenesis in the sea urchin embryo is initiated uniquely in three domains. Development 145, dev167742. 10.1242/dev.167742

70. Meulemans, D., Bronner-Fraser, M., 2007. The Amphioxus *SoxB* Family: Implications for the Evolution of Vertebrate Placodes. Int. J. Biol. Sci. 356–364. 10.7150/ijbs.3.356

71. Minh, B.Q., Schmidt, H.A., Chernomor, O., Schrempf, D., Woodhams, M.D., Von Haeseler, A., Lanfear, R., 2020. IQ-TREE 2: New Models and Efficient Methods for Phylogenetic Inference in the Genomic Era. Mol. Biol. Evol. 37, 1530–1534. 10.1093/molbev/msaa015

72. Mizutani, C.M., Bier, E., 2008. EvoD/Vo: the origins of BMP signalling in the neuroectoderm. Nat. Rev. Genet. 9, 663–677. 10.1038/nrg2417

73. Monjo, F., Romero, R., 2015a. Embryonic development of the nervous system in the planarian Schmidtea polychroa. Dev. Biol. 397, 305–319. 10.1016/j.ydbio.2014.10.021

74. Monjo, F., Romero, R., 2015b. Embryonic development of the nervous system in the planarian Schmidtea polychroa. Dev. Biol. 397, 305–319. 10.1016/j.ydbio.2014.10.021

75. Moore, J.A., Moreno-Campos, R., Noah, A.S., Singleton, E.W., Uribe, R.A., 2025. BMP signaling pathway member expression is enriched in enteric neural progenitors and required for zebrafish enteric nervous system development. Dev. Dyn. 254, 272–287. 10.1002/dvdy.737

76. Mörsdorf, D., Knabl, P., Genikhovich, G., 2024. Highly conserved and extremely evolvable: BMP signalling in secondary axis patterning of Cnidaria and Bilateria. Dev. Genes Evol. 234, 1–19. 10.1007/s00427-024-00714-4

77. Müller, C.H.G., Harzsch, S., Perez, Y., 2018. 7. Chaetognatha, in: Schmidt-Rhaesa, A. (Ed.), Miscellaneous Invertebrates. De Gruyter, pp. 163–282. 10.1515/9783110489279-007

78. Nakanishi, N., Renfer, E., Technau, U., Rentzsch, F., 2012. Nervous systems of the sea anemone *Nematostella vectensis* are generated by ectoderm and endoderm and shaped by distinct mechanisms. Development 139, 347–357. 10.1242/dev.071902

79. Odden, J.P., Holbrook, S., Doe, C.Q., 2002. *Drosophila HB9* Is Expressed in a Subset of Motoneurons and Interneurons, Where It Regulates Gene Expression and Axon Pathfinding. J. Neurosci. 22, 9143–9149. 10.1523/JNEUROSCI.22-21-09143.2002

80. Okuda, Y., Yoda, H., Uchikawa, M., Furutani-Seiki, M., Takeda, H., Kondoh, H., Kamachi, Y., 2006. Comparative genomic and expression analysis of group B1 *sox* genes in zebrafish indicates their diversification during vertebrate evolution. Dev. Dyn. 235, 811–825. 10.1002/dvdy.20678

81. Ordoñez, J.F., Wollesen, T., 2025. Anterior-posterior patterning in the chaetognath Spadella cephaloptera informs bilaterian nervous system and tail evolution. Commun. Biol. 10.1038/s42003-025-09398-6

82. Ordoñez, J.F., Wollesen, T., 2024. Unfolding the ventral nerve center of chaetognaths. Neural Develop. 19, 5. 10.1186/s13064-024-00182-6

83. Overton, P.M., Meadows, L.A., Urban, J., Russell, S., 2002. Evidence for differential and redundant function of the Sox genes *Dichaete* and *SoxN* during CNS development in *Drosophila*. Development 129, 4219–4228. 10.1242/dev.129.18.4219

84. Pasini, A., Amiel, A., Rothbächer, U., Roure, A., Lemaire, P., Darras, S., 2006. Formation of the Ascidian Epidermal Sensory Neurons: Insights into the Origin of the Chordate Peripheral Nervous System. PLoS Biol. 4, e225. 10.1371/journal.pbio.0040225

85. Perez, Y., Rieger, V., Martin, E., Müller, C.H.G., Harzsch, S., 2013. Neurogenesis in an Early Protostome Relative: Progenitor Cells in the Ventral Nerve Center of Chaetognath Hatchlings Are Arranged in a Highly Organized Geometrical Pattern. J. Exp. Zoolog. B Mol. Dev. Evol. 320, 179–193. 10.1002/jez.b.22493

86. Quan, X.-J., Hassan, B.A., 2005. From skin to nerve: flies, vertebrates and the first helix. Cell. Mol. Life Sci. 62, 2036–2049. 10.1007/s00018-005-5124-1

87. Rahman, M.S., Akhtar, N., Jamil, H.M., Banik, R.S., Asaduzzaman, S.M., 2015. TGF-β/BMP signaling and other molecular events: regulation of osteoblastogenesis and bone formation. Bone Res. 3, 15005. 10.1038/boneres.2015.5

88. Rieger, V., Perez, Y., Müller, C.H.G., Lacalli, T., Hansson, B.S., Harzsch, S., 2011. Development of the nervous system in hatchlings of Spadella cephaloptera (Chaetognatha), and implications for nervous system evolution in Bilateria: Nervous system development in S. cephaloptera. Dev. Growth Differ. 53, 740–759. 10.1111/j.1440-169X.2011.01283.x

89. Rieger, V., Perez, Y., Müller, C.H.G., Lipke, E., Sombke, A., Hansson, B.S., Harzsch, S., 2010. Immunohistochemical analysis and 3D reconstruction of the cephalic nervous system in Chaetognatha: insights into the evolution of an early bilaterian brain? Invertebr. Biol. 129, 77–104. 10.1111/j.1744-7410.2010.00189.x

90. Robinow, S., White, K., 1991. Characterization and spatial distribution of the ELAV protein during *Drosophila melanogaster* development. J. Neurobiol. 22, 443–461. 10.1002/neu.480220503

91. Rusten, T.E., Cantera, R., Kafatos, F.C., Barrio, R., 2002. The role of TGFβ signaling in the formation of the dorsal nervous system is conserved between *Drosophila* and chordates. Development 129, 3575–3584. 10.1242/dev.129.15.3575

92. Saha, M.S., Miles, R.R., Grainger, R.M., 1997. Dorsal-Ventral Patterning during Neural Induction inXenopus:Assessment of Spinal Cord Regionalization withxHB9,a Marker for the Motor Neuron Region. Dev. Biol. 187, 209–223. 10.1006/dbio.1997.8625

93. Schindelin, J., Arganda-Carreras, I., Frise, E., Kaynig, V., Longair, M., Pietzsch, T., Preibisch, S., Rueden, C., Saalfeld, S., Schmid, B., Tinevez, J.Y., White, D.J., Hartenstein, V., Eliceiri, K., Tomancak, P., Cardona, A., 2012. Fiji: An open-source platform for biological-image analysis. Nat. Methods. 10.1038/nmeth.2019

94. Schmidt, H., Rickert, C., Bossing, T., Vef, O., Urban, J., Technau, G.M., 1997. The Embryonic Central Nervous System Lineages of Drosophila melanogaster. Dev. Biol. 189, 186–204. 10.1006/dbio.1997.8660

95. Schmidt-Rhaesa, A., 2007. The Evolution of Organ Systems. OUP, Oxford, UK.

96. Shinn, L.G., 1997. Chapter 3: Chaetognatha., in: Harrison, F.W., Ruppert, E.E. (Eds.), Microscopic Anatomy of Invertebrates. Hemichordata, Chaetognatha, and the Invertebrate Chordates. Wiley-Liss, New York, pp. 103–220.

97. Simionato, E., Kerner, P., Dray, N., Le Gouar, M., Ledent, V., Arendt, D., Vervoort, M., 2008. atonal-and achaete-scute-related genes in the annelid Platynereis dumerilii: insights into the evolution of neural basic-Helix-Loop-Helix genes. BMC Evol. Biol. 8, 170. 10.1186/1471-2148-8-170

98. Sloley, B.D., Juorio, A.V., 1995. Monoamine Neurotransmitters in Invertebrates and Vertebrates: An Examination of the Diverse Enzymatic Pathways Utilized to Synthesize and Inactivate Blogenic Amines, in: International Review of Neurobiology. Elsevier, pp. 253–303. 10.1016/S0074-7742(08)60528-0

99. Sprecher, S.G., Reichert, H., 2003. The urbilaterian brain: developmental insights into the evolutionary origin of the brain in insects and vertebrates. Arthropod Struct. Dev. 32, 141–156. 10.1016/S1467-8039(03)00007-0

100. Steenwyk, J.L., Buida, T.J., Li, Y., Shen, X.-X., Rokas, A., 2020. ClipKIT: A multiple sequence alignment trimming software for accurate phylogenomic inference. PLOS Biol. 18, e3001007. 10.1371/journal.pbio.3001007

101. Sur, A., Magie, C.R., Seaver, E.C., Meyer, N.P., 2017. Spatiotemporal regulation of nervous system development in the annelid Capitella teleta. EvoDevo 8, 13. 10.1186/s13227-017-0076-8

102. Sur, A., Renfro, A., Bergmann, P.J., Meyer, N.P., 2020. Investigating cellular and molecular mechanisms of neurogenesis in Capitella teleta sheds light on the ancestor of Annelida. BMC Evol. Biol. 20, 84. 10.1186/s12862-020-01636-1

103. Tan, S., Huan, P., Liu, B., 2022. Molluskan Dorsal–Ventral Patterning Relying on BMP2/4 and Chordin Provides Insights into Spiralian Development and Evolution. Mol. Biol. Evol. 39, msab322. 10.1093/molbev/msab322

104. Vervoort, M., Ledent, V., 2001. The Evolution of the Neural Basic Helix-Loop-Helix Proteins. Sci. World J. 1, 396–426. 10.1100/tsw.2001.68

105. Vidal, B., Santella, A., Serrano-Saiz, E., Bao, Z., Chuang, C.-F., Hobert, O., 2015. *C. elegans* SoxB genes are dispensable for embryonic neurogenesis but required for terminal differentiation of specific neuron types. Development dev.125740. 10.1242/dev.125740

106. Waki, K., Imai, K.S., Satou, Y., 2015. Genetic pathways for differentiation of the peripheral nervous system in ascidians. Nat. Commun. 6, 8719. 10.1038/ncomms9719

107. Wegner, M., Stolt, C.C., 2005. From stem cells to neurons and glia: a Soxist’s view of neural development. Trends Neurosci. 28, 583–588. 10.1016/j.tins.2005.08.008

108. Wollesen, T., Rodriguez Monje, S.V., Oel, A.P., Arendt, D., 2023a. Characterization of eyes, photoreceptors, and opsins in developmental stages of the arrow worm *Spadella cephaloptera* (Chaetognatha). J. Exp. Zoolog. B Mol. Dev. Evol. 340, 342–353. 10.1002/jez.b.23193

109. Wollesen, T., Rodriguez Monje, S. V, Oel, A.P., Arendt, D., 2023b. Characterization of eyes, photoreceptors, and opsins in developmental stages of the arrow worm Spadella cephaloptera (Chaetognatha). J. Exp. Zoolog. B Mol. Dev. Evol. 10.1002/jez.b.23193

110. Zakin, L., De Robertis, E.M., 2010. Extracellular regulation of BMP signaling. Curr. Biol. CB 20, R89–92. 10.1016/j.cub.2009.11.021

111. Zhao, G., Skeath, J.B., 2002. The Sox-domain containing gene *Dichaete/fish-hook* acts in concert with *vnd* and *ind* to regulate cell fate in the *Drosophila* neuroectoderm. Development 129, 1165–1174. 10.1242/dev.129.5.1165

112. Zhou, Z., Wang, L., Yang, J., Zhang, H., Kong, P., Wang, M., Qiu, L., Song, L., 2011. A dopamine beta hydroxylase from Chlamys farreri and its induced mRNA expression in the haemocytes after LPS stimulation. Fish Shellfish Immunol. 30, 154–162. 10.1016/j.fsi.2010.09.020

