## Supplementary Figure 1 - 13 for "Early nervous system development in the chaetognath *Spadella cephaloptera* exhibits conserved bilaterian patterning features"

*Corresponding authors

Tim Wollesen

June F. Ordoñez

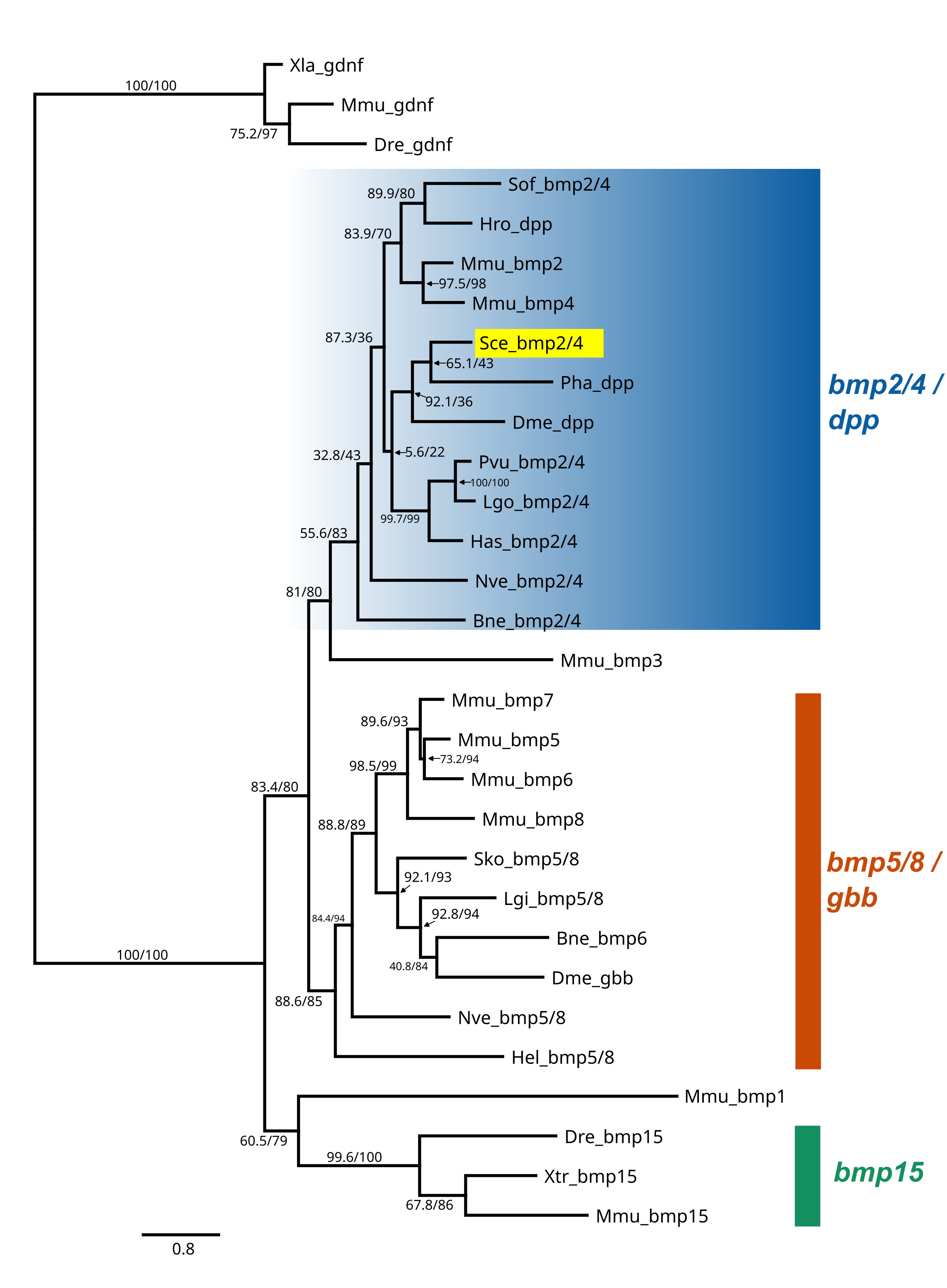

**Supplementary Figure 1.** Maximum Likelihood phylogenetic analysis of bone morphogenetic protein (BMP) amino acid sequences including the deduced Bmp2/4/Dpp protein from *Spadella cephaloptera* (highlighted in yellow). The tree was inferred using IQ-TREE based on bilaterian protein sequences obtained from published sources and NCBI GenBank BLAST searches. Branch support values are shown as SH-aLRT (%) / ultrafast bootstrap support (UFBS %). Species abbreviations are provided in Supplementary Table 1.

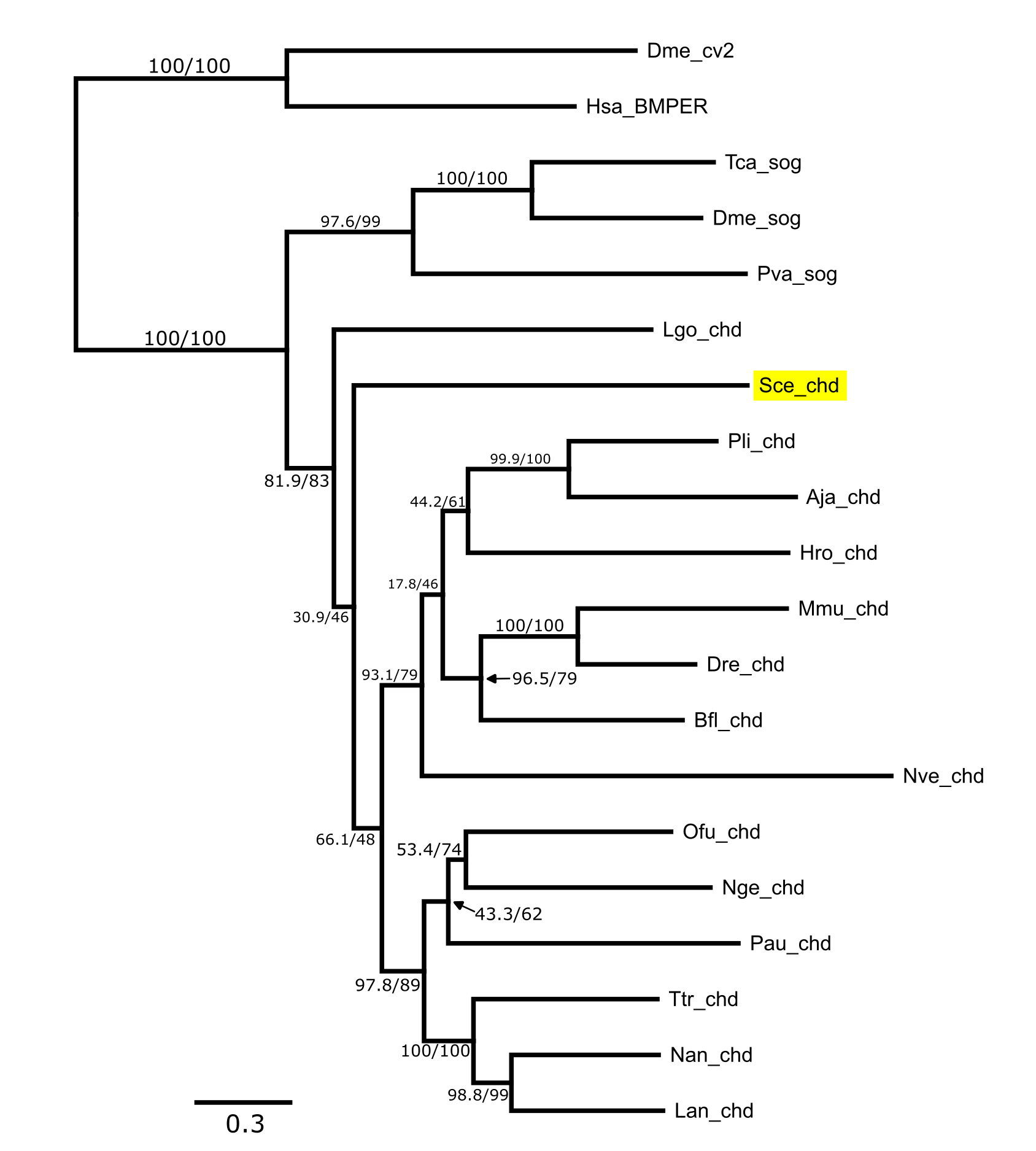

**Supplementary Figure 2.** Maximum Likelihood phylogenetic analysis of Chordin/Short gastrulation (Sog) amino acid sequences including deduced Chd/Sog protein from *Spadella cephaloptera* (highlighted in yellow). The tree was inferred using IQ-TREE based on bilaterian protein sequences obtained from published sources and NCBI GenBank BLAST searches. Branch support values are shown as SH-aLRT (%) / ultrafast bootstrap support (UFBS %). Species abbreviations are provided in Supplementary Table 1.

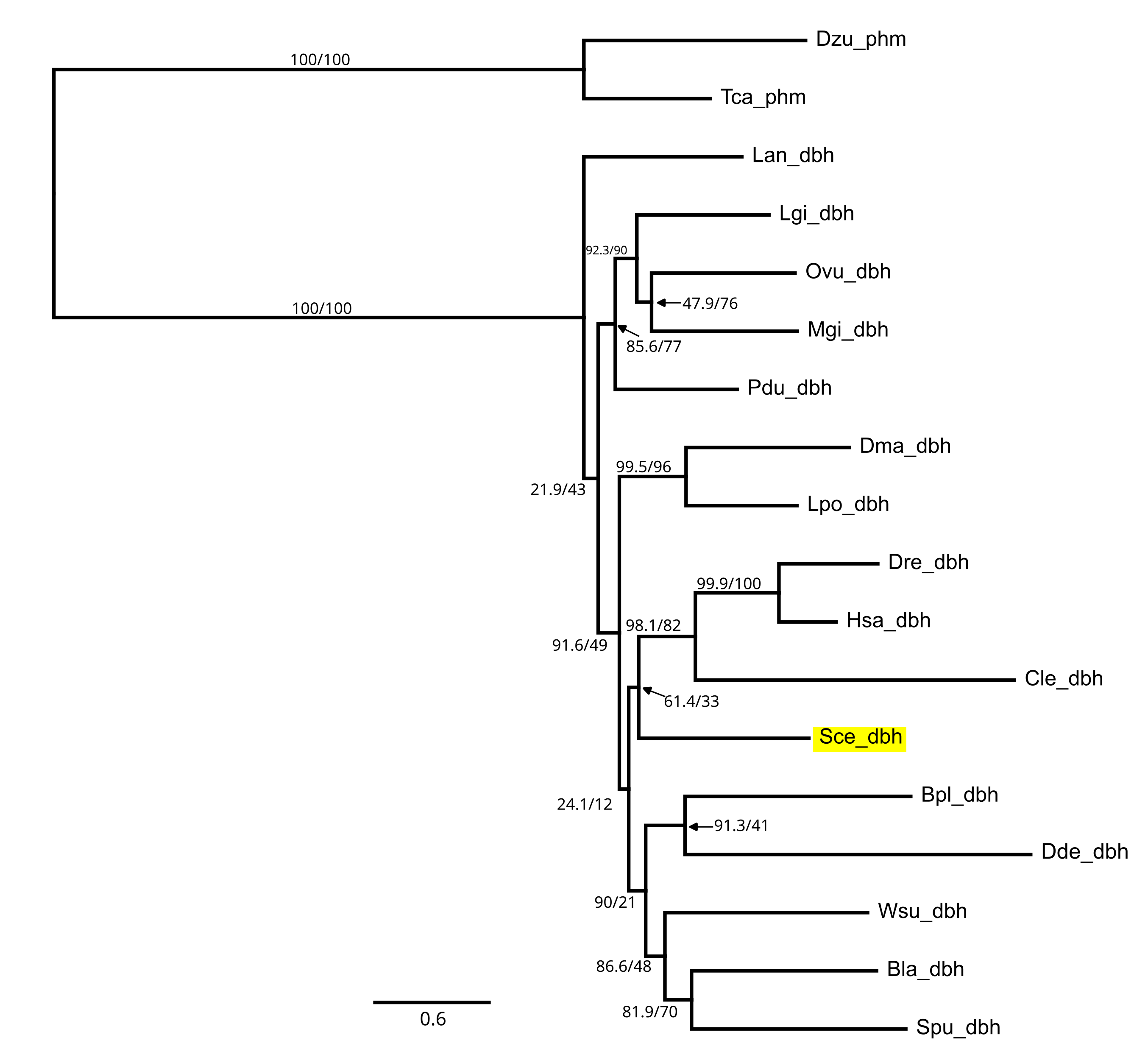

**Supplementary Figure 3.** Maximum Likelihood phylogenetic analysis of Dopamine-β-hydroxylase (Dbh) amino acid sequences including deduced Dbh protein from *Spadella cephaloptera* (highlighted in yellow). The tree was inferred using IQ-TREE based on bilaterian protein sequences obtained from published sources and NCBI GenBank BLAST searches. Branch support values are shown as SH-aLRT (%) / ultrafast bootstrap support (UFBS %). Species abbreviations are provided in Supplementary Table 1.

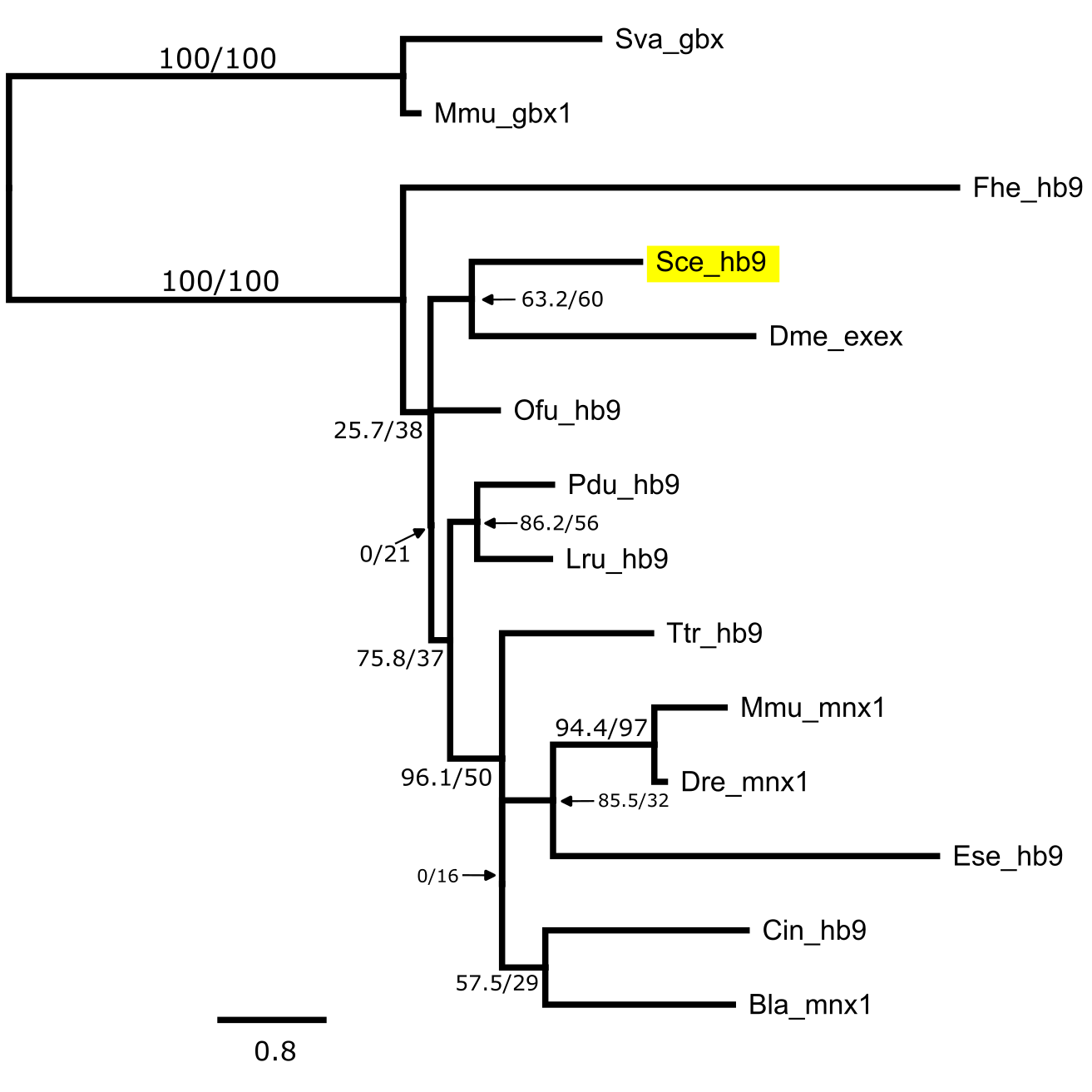

**Supplementary Figure 4.** Maximum Likelihood phylogenetic analysis of Hb9/Motor neuron and pancreas homeobox 1 (Mnx1)/Extra-extra (ex-ex) amino acid sequences including deduced Hb9/Mnx1/Exex protein from *Spadella cephaloptera* (highlighted in yellow). The tree was inferred using IQ-TREE based on bilaterian protein sequences obtained from published sources and NCBI GenBank BLAST searches. Branch support values are shown as SH-aLRT (%) / ultrafast bootstrap support (UFBS %). Species abbreviations are provided in Supplementary Table 1.

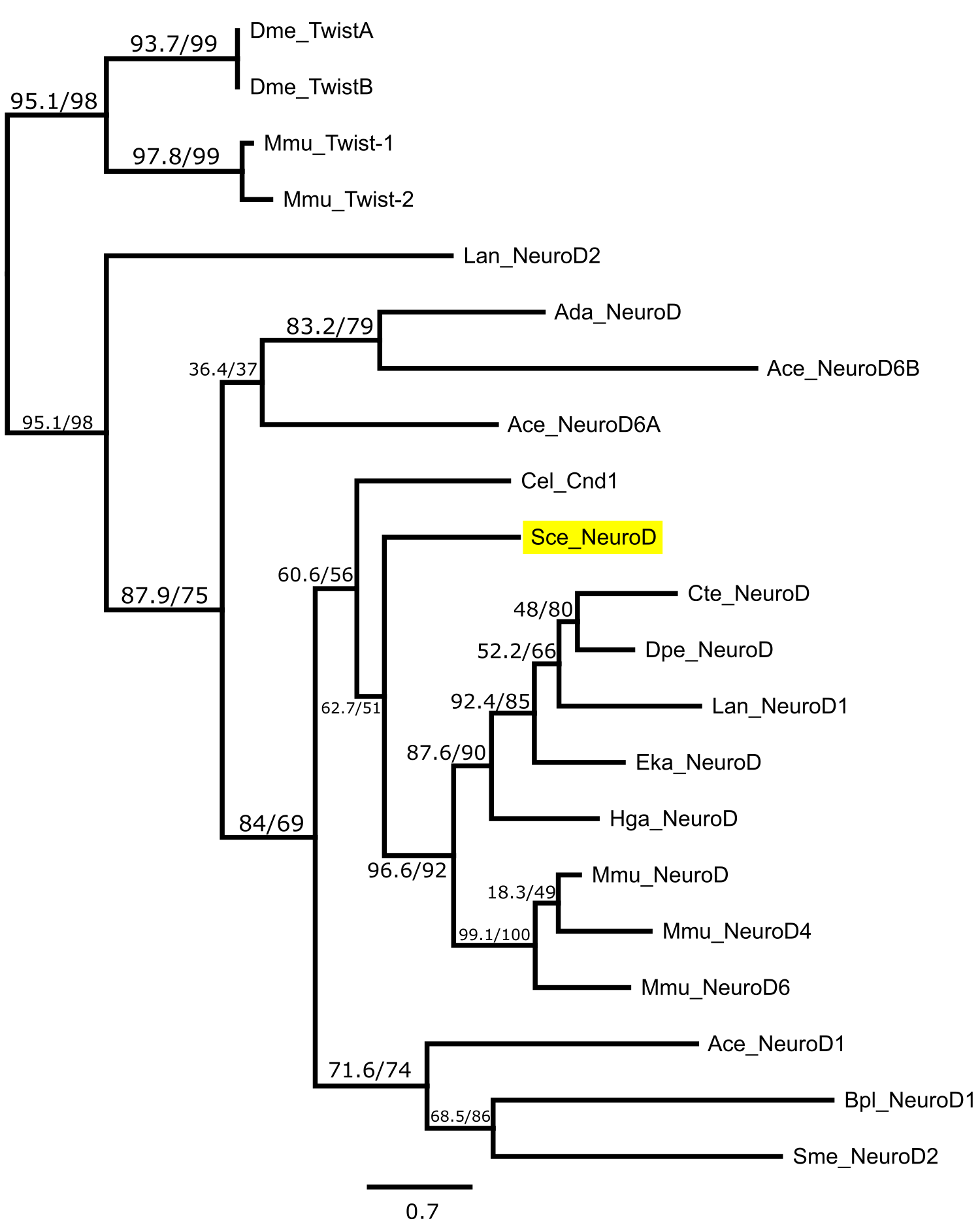

**Supplementary Figure 5.** Maximum Likelihood phylogenetic analysis of Neuronal differentiation 1 (NeuroD) amino acid sequences including deduced NeuroD protein from *Spadella cephaloptera* (highlighted in yellow). The tree was inferred using IQ-TREE based on bilaterian protein sequences obtained from published sources and NCBI GenBank BLAST searches. Branch support values are shown as SH-aLRT (%) / ultrafast bootstrap support (UFBS %). Species abbreviations are provided in Supplementary Table 1.

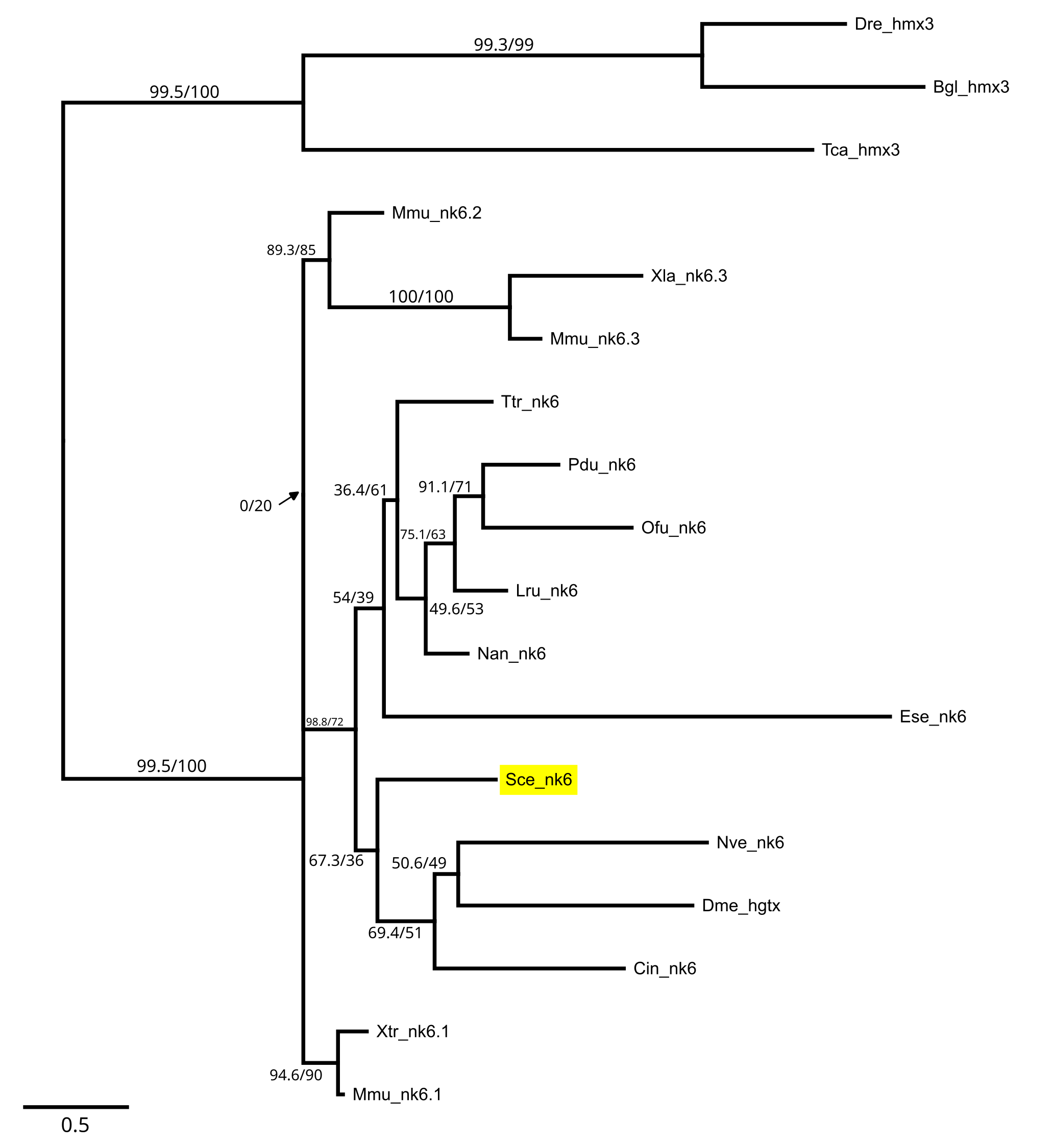

**Supplementary Figure 6.** Maximum Likelihood phylogenetic analysis of Nk6/Hgtx amino acid sequences including deduced Nk6/Hgtx protein from *Spadella cephaloptera* (highlighted in yellow). The tree was inferred using IQ-TREE based on bilaterian protein sequences obtained from published sources and NCBI GenBank BLAST searches. Branch support values are shown as SH-aLRT (%) / ultrafast bootstrap support (UFBS %). Species abbreviations are provided in Supplementary Table 1.

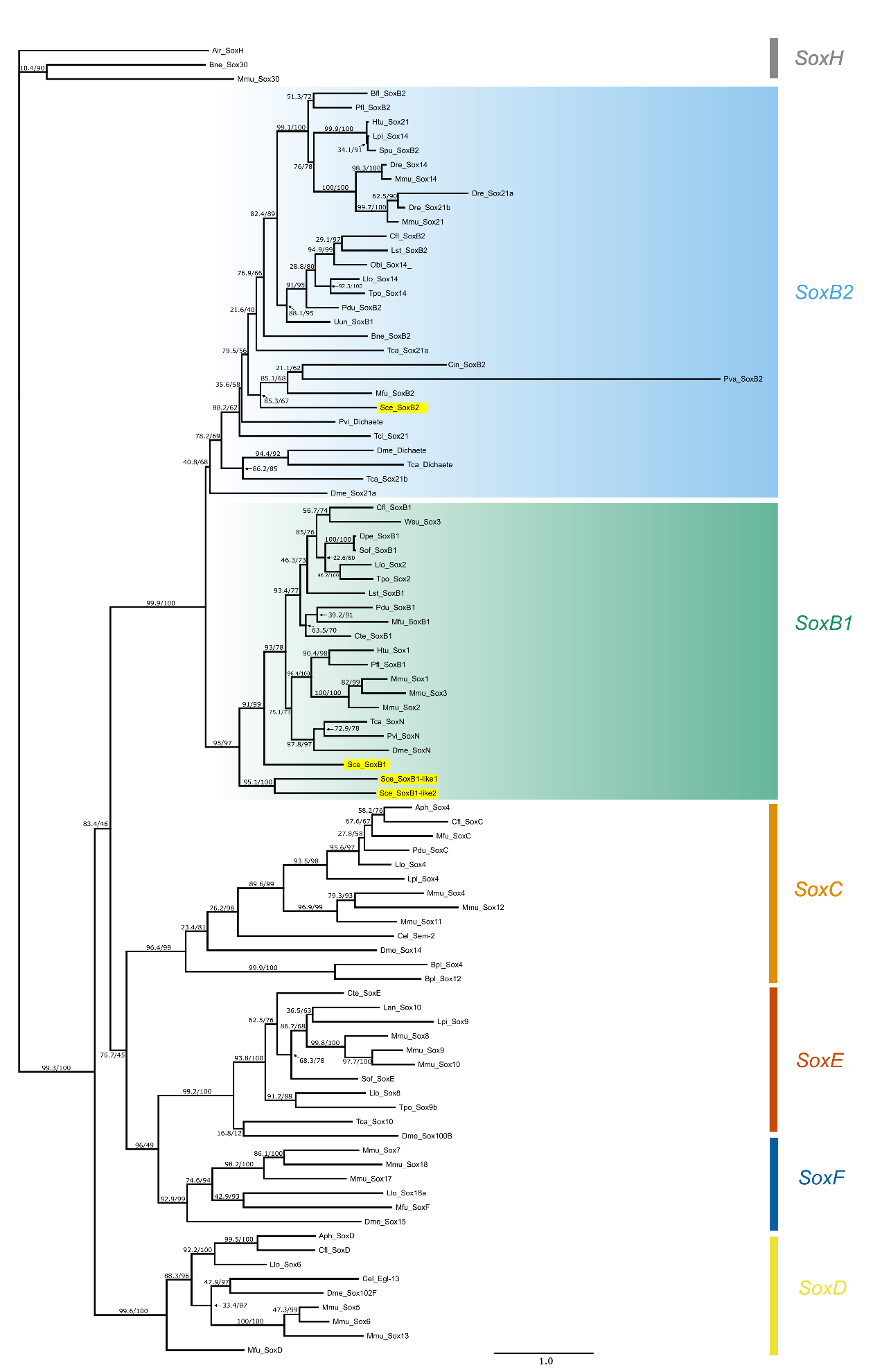

**Supplementary Figure 7.** Maximum Likelihood phylogenetic analysis of SRY-related HMG box (Sox) amino acid sequences including deduced protein SoxB protein from *Spadella cephaloptera* (highlighted in yellow). The unrooted tree was inferred using IQ-TREE based on bilaterian protein sequences obtained from published sources and NCBI GenBank BLAST searches. Branch support values are shown as SH-aLRT (%) / ultrafast bootstrap support (UFBS %). SoxB1 and SoxB2 clades are highlighted in dark teal and light blue, respectively. Species abbreviations are provided in Supplementary Table 1.

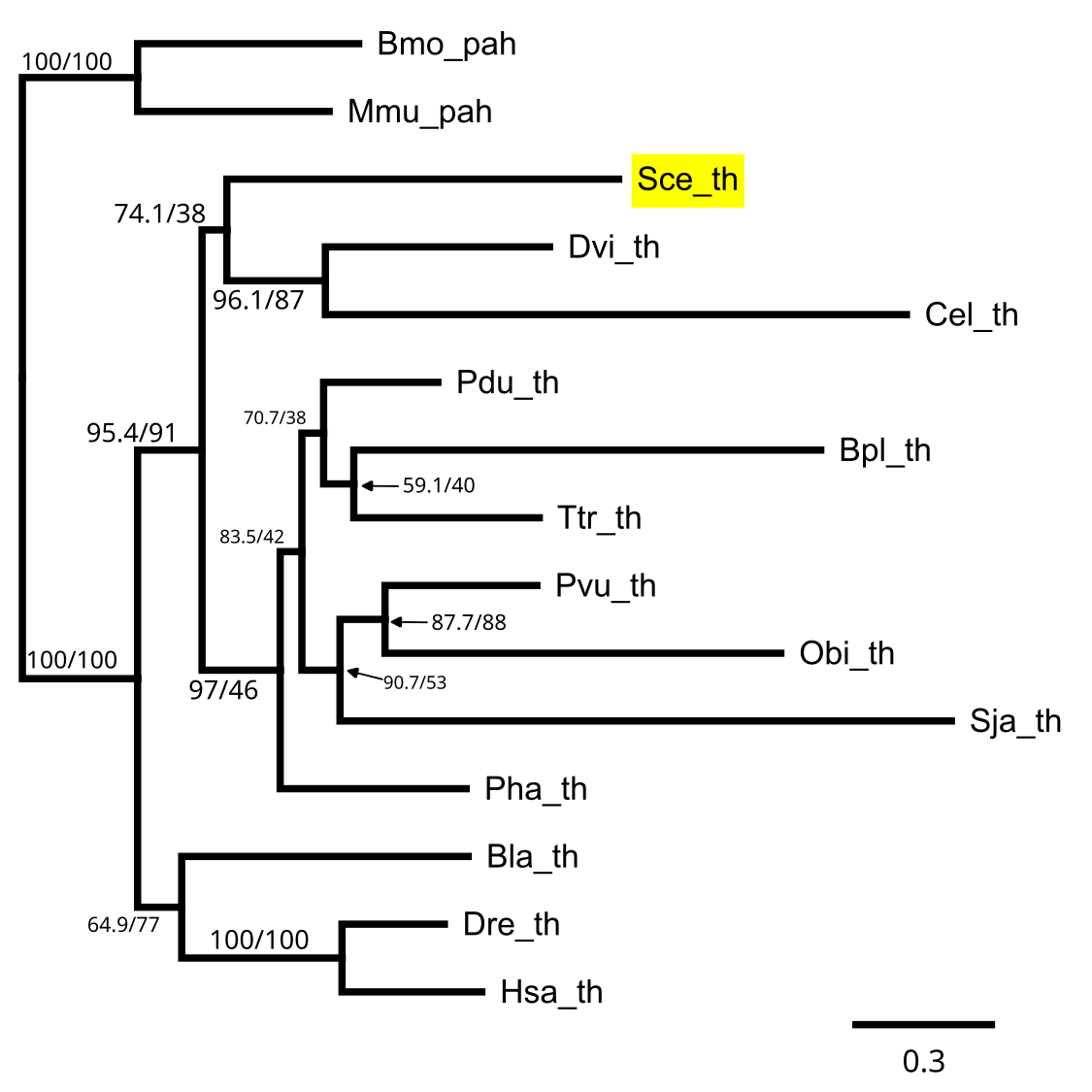

**Supplementary Figure 8.** Maximum Likelihood phylogenetic analysis of Tyrosine Hydroxylase (Th) amino acid sequences including deduced Th protein from *Spadella cephaloptera* (highlighted in yellow). The tree was inferred using IQ-TREE based on bilaterian protein sequences obtained from published sources and NCBI GenBank BLAST searches. Branch support values are shown as SH-aLRT (%) / ultrafast bootstrap support (UFBS %). Species abbreviations are provided in Supplementary Table 1.

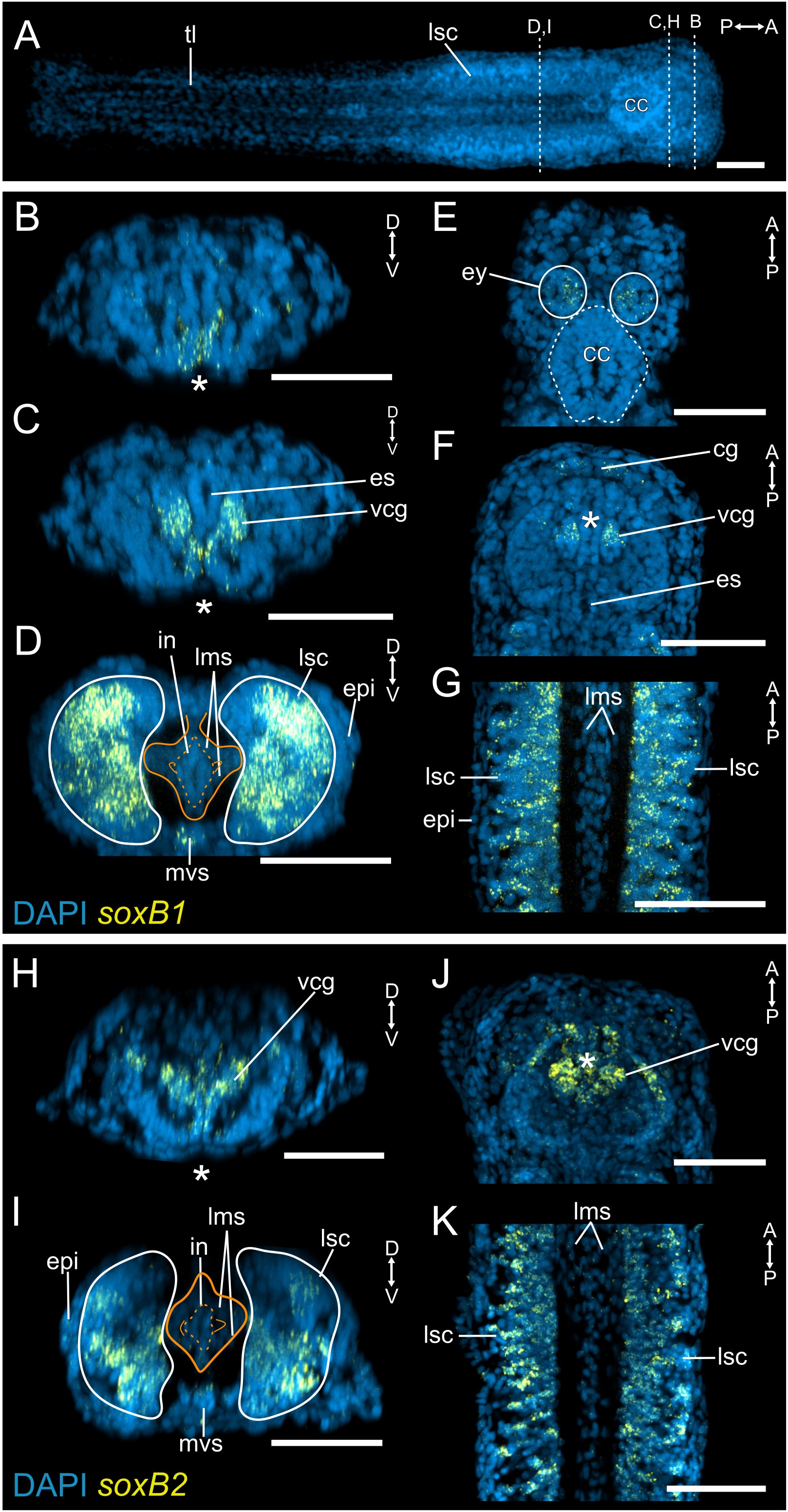

### Supplementary Figure 9. Expression patterns of *Sce-soxB1* and *Sce-soxB2* in the *Spadella cephaloptera* hatchling.

(A) Dorsal maximum projection of a DAPI-counterstained hatchling indicating the positions along the anterior–posterior axis from which transverse sections were collected.

(B–G) *Sce-soxB1* expression. (B–D) Transverse sections through the head (B, C) and trunk (D). Orange outline demarcates the mesodermal derivatives, including the trunk longitudinal muscles. (E, F) Dorsal sections of the head. (E) dorsal-most section showing expression in the eyes. (F) mid-level section showing expression in the presumptive anteroventral cephalic ganglion anlage. (G) Mid-dorsal section of the trunk.

(H–J) *Sce-soxB2* expression. (H, I) Transverse sections through the head (H) and trunk (I). (J) Mid-dorsal sections of the head and trunk.

Scale bars: 50 µm, except (A): 100 µm. Asterisks indicate the position of the future mouth opening. Orientation is indicated in the upper right corner of each panel. Abbreviations: cc, corona ciliata; epi, epidermis; ey, eye; in, intestine; lsc, lateral somata clusters; lms, longitudinal muscle somata; mvs, medioventral somata clusters; vcg, presumptive anteroventral cephalic ganglion anlage; tl: tail.

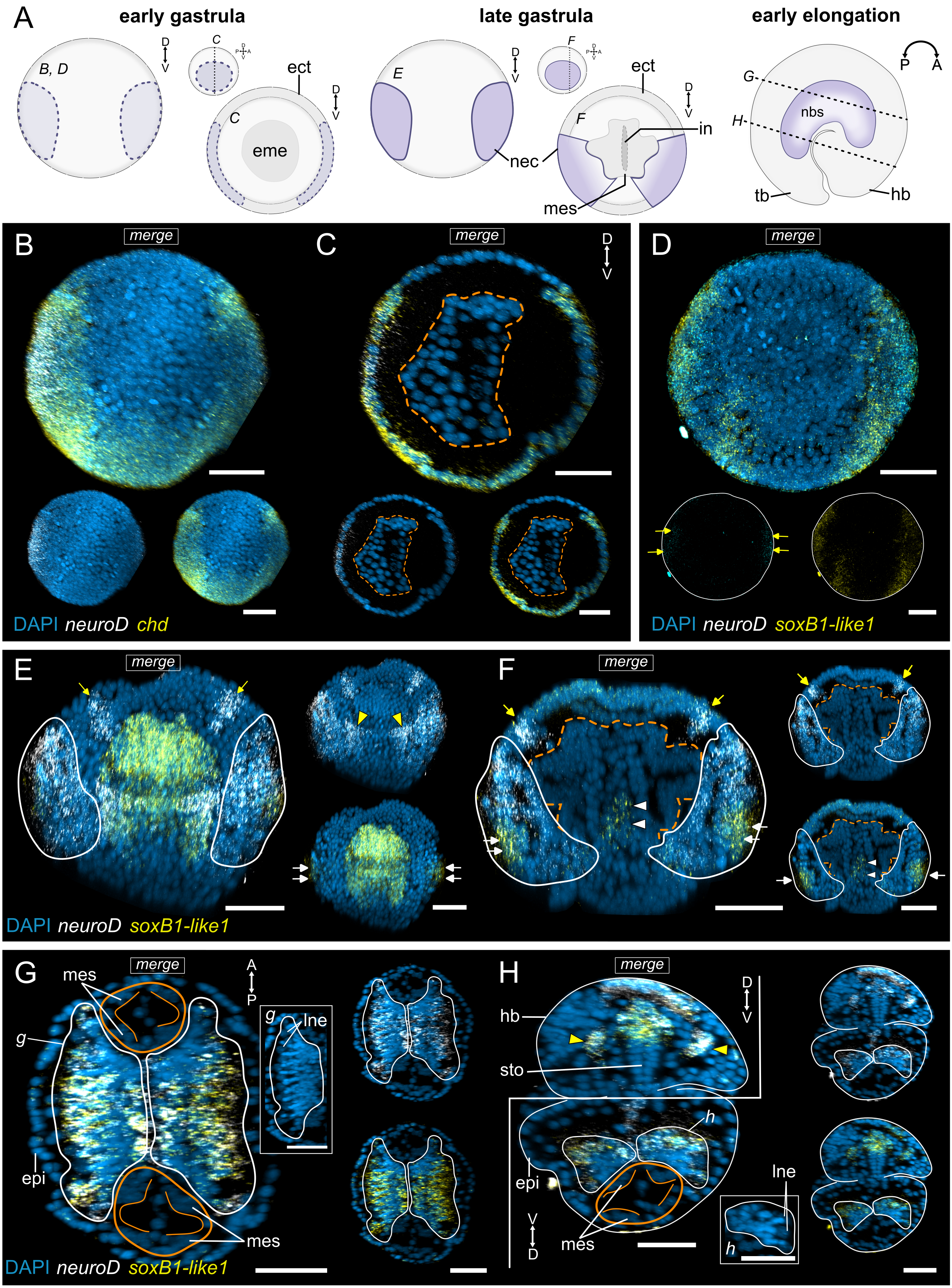

### Supplementary Figure 10. Double-label expression analyses of *Sce-neuroD* with *Sce-soxB1-like1* and *Sce-chd* during embryonic development of *Spadella cephaloptera*. Fluorescence panels are shown as a composite overlay (DAPI and two probe signals) together with the corresponding individual probe channels shown with DAPI counterstain.

(A) Schematic representations of early gastrula, late gastrula, and early elongation stages. Early and late gastrula stages are shown in transverse views (full view and transverse section), while the early elongation stage is shown in lateral view. Neurogenic regions, including the NEC and neural cells of the nascent VNC, are indicated in purple.

(B–D) Early gastrula. (B) Transverse maximum projection and (C) transverse section showing *Sce-neuroD* and *Sce-chordin* expression. Orange dashed outline demarcates the endomesoderm. (D) Transverse maximum projection showing *Sce-neuroD* and *Sce-soxB1-like1* expression. Separate channels are shown without DAPI to improve visualization. Yellow arrows indicate *Sce-neuroD* expression domains.

(E, F) Late gastrula. (E) Transverse maximum projection and (F) transverse section. Yellow arrows indicate *Sce-neuroD* expression detected outside the anatomically defined NEC; yellow arrowheads highlight paired anterior ectodermal expression domains; white arrows mark the *Sce-soxB1-like1* expression in the ventral NEC; white arrowheads marks *soxB1-like1* expression in the anterior endomesoderm. White outlines demarcate the NEC.

(G, H) Early elongation. (G) Dorsal and (H) transverse section through the early elongation trunk. Orange outlines demarcate the mesodermal cells. Yellow arrows highlight paired *Sce-neuroD*⁺/*Sce-soxB1-like1*⁺ expression domains in the head bud. Insets *g* and *h* show the corresponding DAPI-only views of the left portion of the nascent VNC.

Scale bars: 50 µm. Orientation is indicated in the schematic representations and, where applicable, in the upper right corner of each panel. Abbreviations: ect, ectoderm; eme, endomesoderm; epi, epidermis; hb, head bud; in, intestine; lne, large-nucleus neuroectodermal cells; mes, mesodermal cells; nbs, neural cells of the developing VNC; nec, neuroectoderm; sto, stomodeum; tb, tail bud.

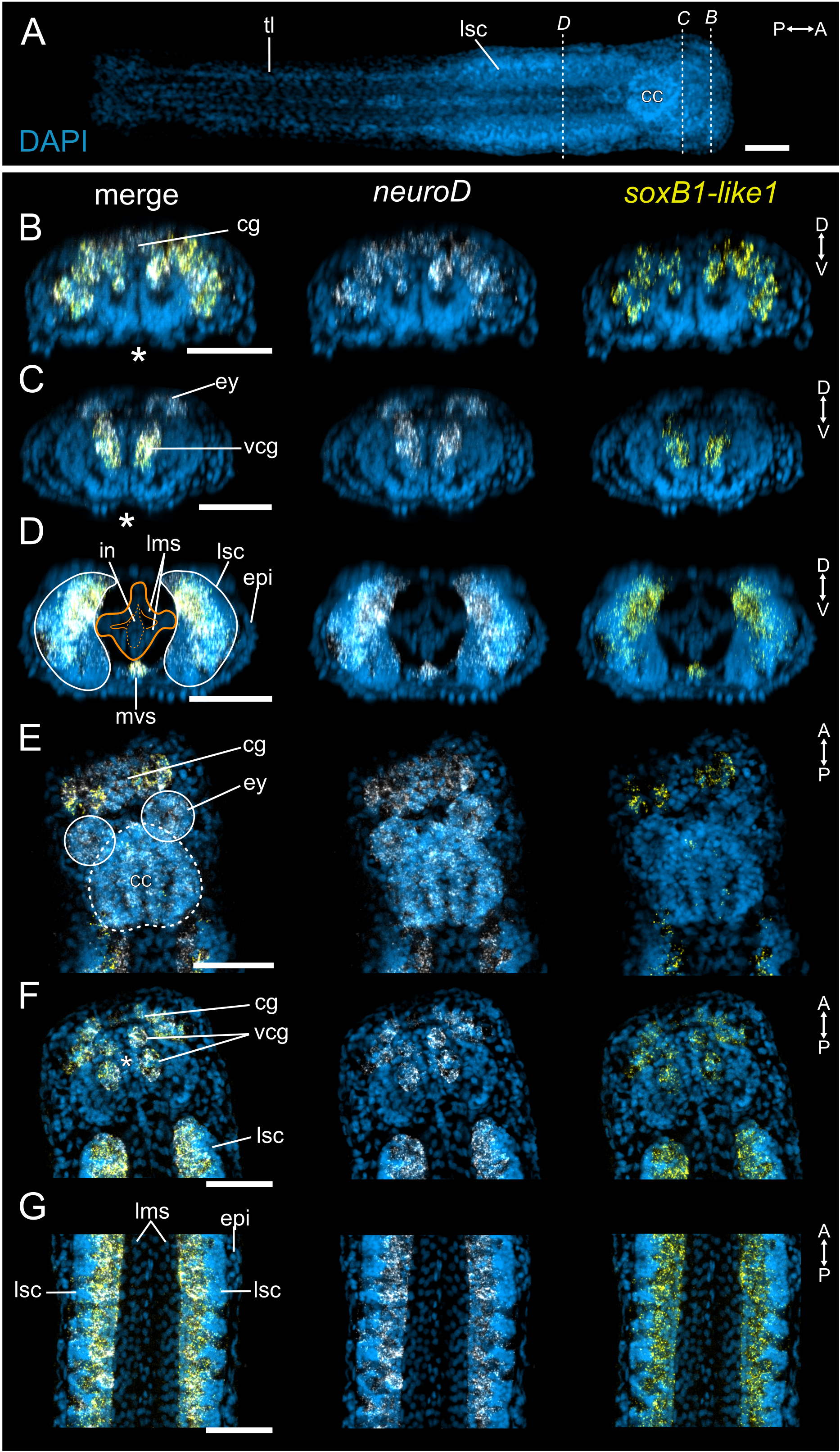

### Supplementary Figure 11. Double-label expression analyses of *Sce-neuroD* and *Sce-soxB1-like1* in the *Spadella cephaloptera* hatchling. Fluorescence panels are shown as a composite overlay (DAPI and two probe signals) together with the corresponding individual probe channels shown with DAPI counterstain.

(A) Dorsal maximum projection of a DAPI-counterstained hatchling indicating the positions along the anterior–posterior axis from which transverse sections were collected.

(B–G) Expression shown as merged and separate channels. (B–D) Transverse sections through the head (B, C) and trunk (D). Orange outline demarcates the mesodermal derivatives, including the trunk longitudinal muscles, and the intestine (dashed). (E, F) Dorsal sections of the head. (E) Dorsal-most section showing expression of *Sce-neuroD* in the eyes and corona ciliata. (F) Mid-level section showing expression in the presumptive anteroventral cephalic ganglion anlage and cephalic ganglion. (G) Mid-dorsal section of the trunk.

Scale bars: 50 µm, except (A): 100 µm. Asterisks indicate the position of the future mouth opening. Orientation is indicated in the upper right corner of each panel. Abbreviations: cc, corona ciliata; cg, cerebral ganglion; epi, epidermis; ey, eye; in, intestine; lsc, lateral somata clusters; lms, longitudinal muscle somata; mvs, medioventral somata clusters; vcg, presumptive anteroventral cephalic ganglion anlage; tl: tail.

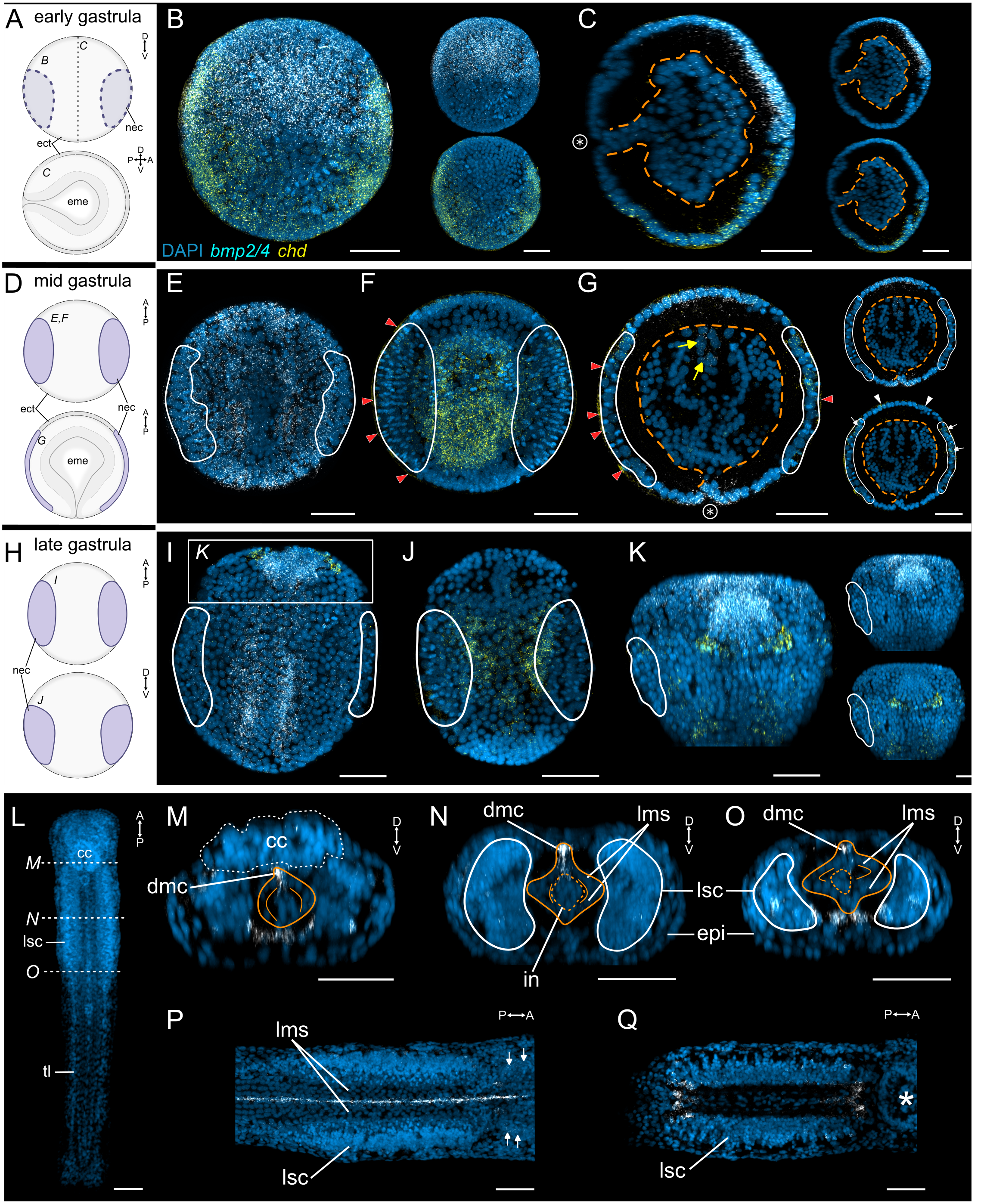

### Supplementary Figure 12. Double-label expression analyses of *Sce-bmp2/4* and *Sce-chd* during embryonic development and *Sce-bmp2/4* expression in the hatchling of *Spadella cephaloptera*. Where applicable, fluorescence panels are shown as a composite overlay (DAPI and two probe signals) together with the corresponding individual probe channels shown with DAPI counterstain.

(A, D, H) Schematic representations of early (A), mid (D), and late (H) gastrula stages. The early gastrula is shown in transverse view (top) and as a lateral section (bottom); the mid gastrula is shown in full dorsal view (top) and as a dorsal section (bottom); the late gastrula is shown in full dorsal view (top) and full transverse view (bottom). The NEC is indicated in purple.

(B, C) Early gastrula. (B) Transverse maximum projection and (C) lateral section through the blastoporal region showing merged channels (left) and individual channels (right). Dashed outlines demarcate the endomesoderm. Encircled asterisks mark the blastoporal region.

(E–G) Mid gastrula. (E, F) Dorsal maximum projections of the dorsal (E) and ventral (F) portions of the embryo. White outlines demarcate the NEC. Red arrowheads indicate non-specific signal from the inner shell layer. (G) Dorsal section through the blastoporal region showing *Sce-bmp2/4* expression. Yellow arrows mark *Sce-bmp2/4* expression in a subset of endomesodermal cells. White arrows indicate *Sce-chd* expression in NEC cells, and white arrowheads highlight anterior ectodermal *Sce-chd* expression.

(I–K) Late gastrula. (I, K) Dorsal maximum projections of the dorsal (I) and ventral (J) portions of the embryo. (K) Transverse maximum projection of the anterior region.

(L–Q) *Sce-bmp2/4* expression in the hatchling. (L) Dorsal maximum projection of a DAPI-counterstained hatchling indicating the positions along the anterior–posterior axis from which transverse sections were collected. (M–O) Transverse sections through the head (M) and trunk (N, O). Orange outlines demarcate mesodermal derivatives, including dorsal medial cells and trunk longitudinal muscles, and the intestine (dashed outline). (P, Q) Dorsal sections of the trunk. (P) Mid-dorsal section showing expression in the dorsal medial cells (yellow arrows) along the trunk and weak signal near the grasping spine region (white arrows). (Q) Ventral section showing expression in cells near the head–trunk and trunk–tail boundaries. Asterisks indicate the position of the future mouth opening.

Scale bars: 50 µm; except (L): 100 µm. Orientation is indicated in the schematic representations and, where applicable, in the upper right corner of each panel. Abbreviations: cc, corona ciliata; ect, ectoderm; eme, endomesoderm; epi, epidermis; dmc, dorsal medial cells; in, intestine; lms, longitudinal muscle somata; nec, neuroectoderm; tl: tail.

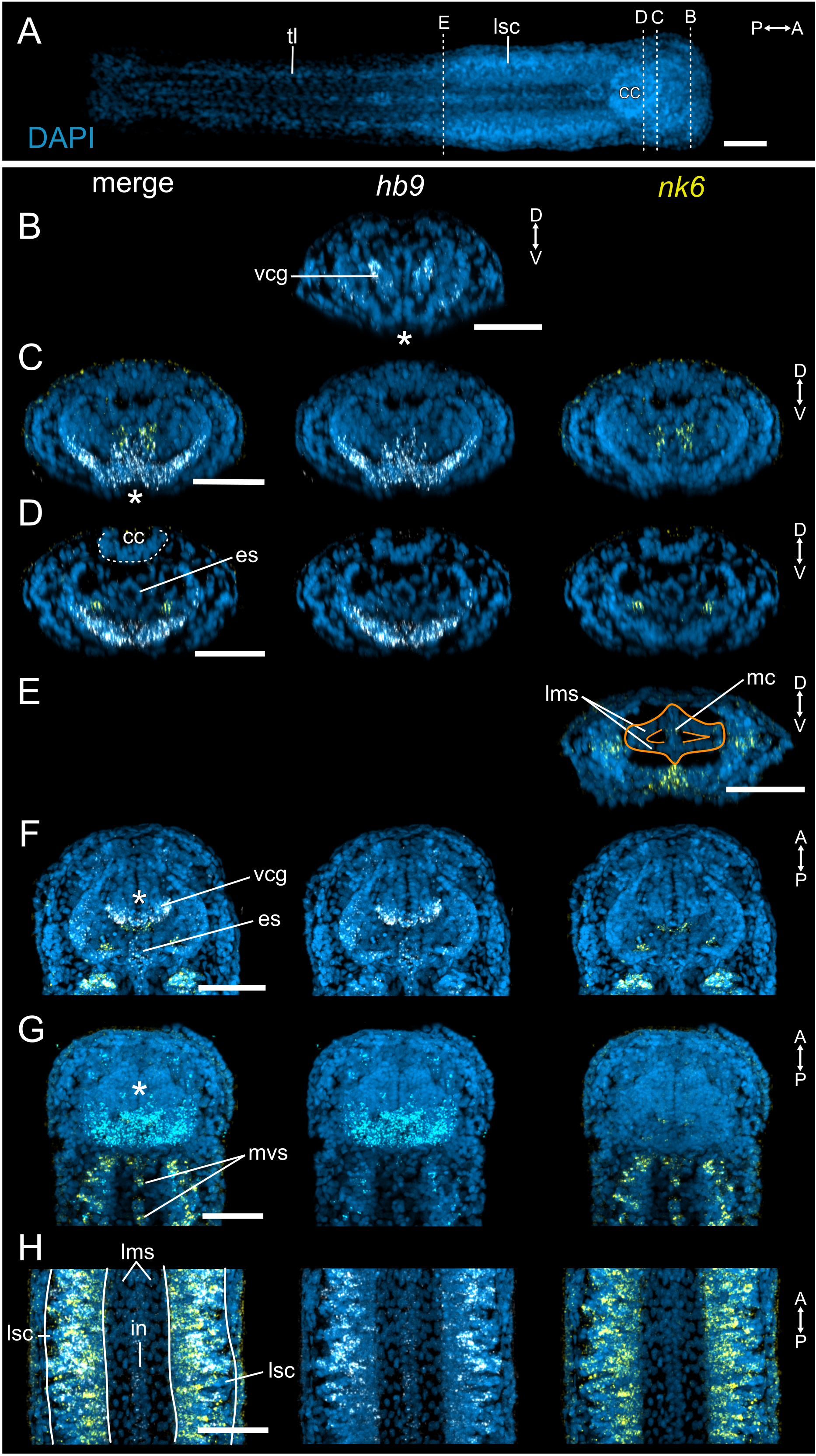

### Supplementary Figure 13. Double-label expression analyses of *Sce-hb9* and *Sce-nk6* in the *Spadella cephaloptera* hatchling. Where applicable, fluorescence panels are shown as a composite overlay (DAPI and two probe signals) together with the corresponding individual probe channels shown with DAPI counterstain.

(A) Dorsal maximum projection of a DAPI-counterstained hatchling indicating the positions along the anterior–posterior axis from which transverse sections were collected.

(B–H) Expression shown as merged images and separate channels.
(B–E) Transverse sections through the head (B–D) and trunk–tail boundary (E). *Sce-hb9* is broadly expressed in ventral head cells corresponding to the presumptive perioral epidermis (B–D), whereas *Sce-nk6* expression is detected in discrete cells of the head (C, D) and posterior trunk (E). Orange outlines demarcate mesodermal derivatives, including mesenterial cells and trunk longitudinal muscles.

(F–H) Dorsal sections. (F) Mid-dorsal section of the head showing expression in the presumptive anteroventral cephalic ganglion anlage and esophagus. (G) Ventral-most section of the head showing *Sce-nk6* expression in the medioventral somata clusters. (H) Mid-dorsal section of the trunk.

Scale bars: 50 µm, except (A): 100 µm. Asterisks indicate the position of the future mouth opening. Orientation is indicated in the upper right corner of each panel. Abbreviations: cc, corona ciliata; epi, epidermis; es, esophagus; in, intestine; lsc, lateral somata clusters; lms, longitudinal muscle somata; mc: mesenterial cells; mvs, medioventral somata clusters; vcg, presumptive anteroventral cephalic ganglion anlage.
